# Archaeogenetic analysis of Neolithic sheep from Anatolia suggests a complex demographic history since domestication

**DOI:** 10.1101/2020.04.17.033415

**Authors:** Erinç Yurtman, Onur Özer, Eren Yüncü, Nihan Dilşad Dağtaş, Dilek Koptekin, Yasin Gökhan Çakan, Mustafa Özkan, Ali Akbaba, Damla Kaptan, Gözde Atağ, Kıvılcım Başak Vural, Can Yümni Gündem, Louise Martin, Gülşah Merve Kılınç, Ayshin Ghalichi, Sinan Can Açan, Reyhan Yaka, Ekin Sağlıcan, Vendela Kempe Lagerholm, Maja Krzewinska, Evangelia Pişkin, Müge Şevketoğlu, C. Can Bilgin, Çiğdem Atakuman, Yılmaz Selim Erdal, Elif Sürer, Johannes Lenstra, Sevgi Yorulmaz, Foad Abazari, Javad Hoseinzadeh, Douglas Baird, Erhan Bıçakçı, Özlem Çevik, Fokke Gerritsen, Rana Özbal, Anders Götherström, Mehmet Somel, İnci Togan, Füsun Özer

## Abstract

Sheep was among the first domesticated animals, but its demographic history is little understood. Here we present combined analyses of mitochondrial and nuclear polymorphism data from ancient central and west Anatolian sheep dating to the Late Glacial and early Holocene. We observe loss of mitochondrial haplotype diversity around 7500 BCE during the early Neolithic, consistent with a domestication-related bottleneck. Post-7000 BCE, mitochondrial haplogroup diversity increases, compatible with admixture from other domestication centres and/or from wild populations. Analysing archaeogenomic data, we further find that Anatolian Neolithic sheep (ANS) are genetically closest to present-day European breeds, and especially those from central and north Europe. Our results indicate that Asian contribution to south European breeds in the post-Neolithic era, possibly during the Bronze Age, may explain this pattern.

## Introduction

Domestication of animals during the Neolithic transition in SW Asia and their spread into new regions had immense economic, demographic, and socio-cultural impacts on human societies^1,2^. Sheep was one of the four main animal species managed and domesticated in this process. Archaeological evidence indicates that sedentary human communities were practicing sheep management already by 9,000-8,000 BCE in an area ranging from central Turkey to northwest Iran^3,4^; this is evidenced, for instance, by signs of corralling in the central Anatolian site Asikli Höyük^5–7^ and young male kill-off practices identified in southeast Anatolian Çayönü and Nevali Çori^8,9^ (Fig. 1). After 7500 BCE, young male kill-off as well as domestication-related morphological changes, such as small size, became widespread across the Fertile Crescent, as in the 7th millennium central Anatolian site of Çatalhöyük^9^. Following 7000 BCE, along with other elements of Neolithic lifeways, humans spread domesticated sheep to neighbouring regions, including Europe, north Africa, and central Asia^3,4^.

**Fig. 1.**
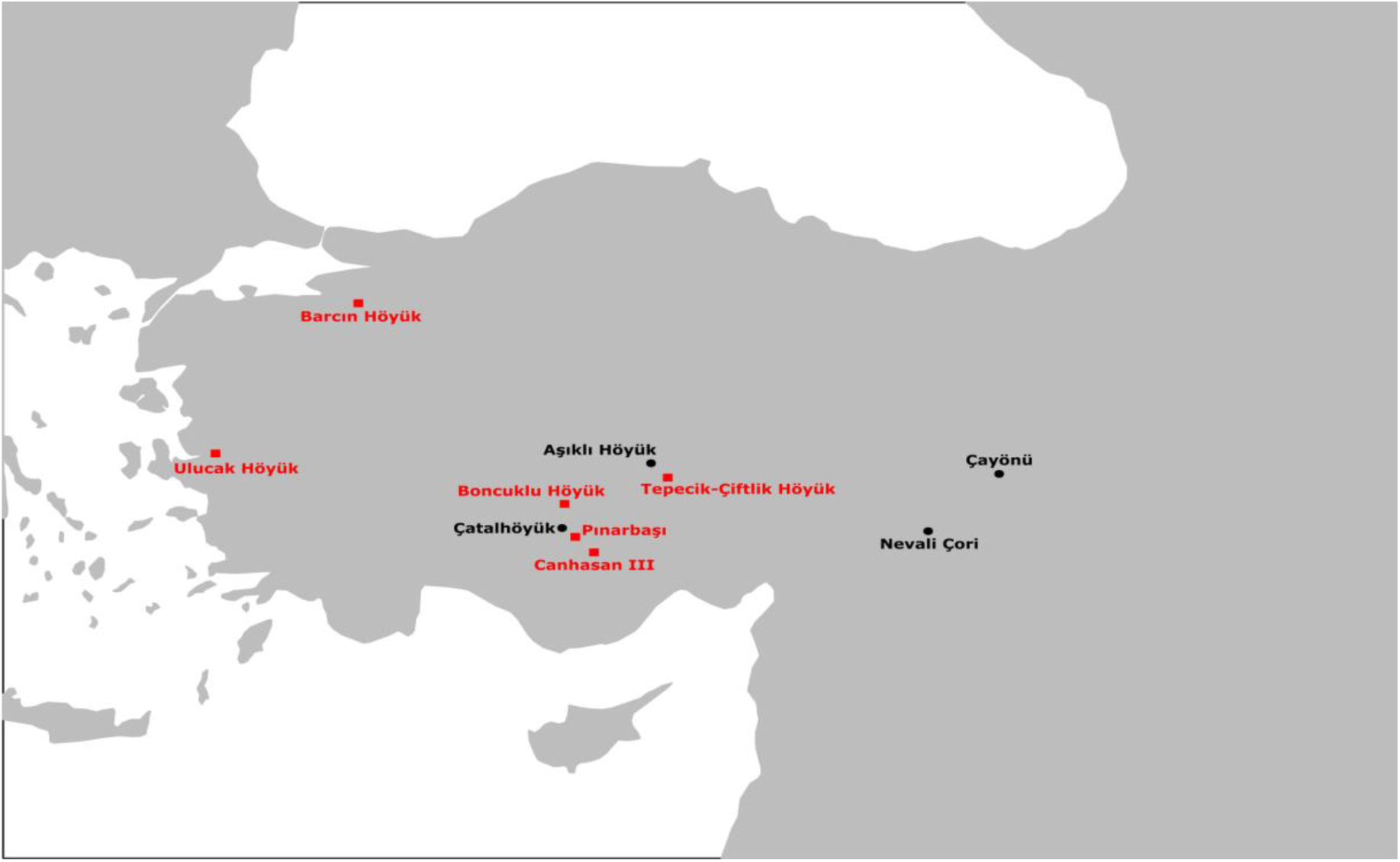
Map of archaeological sites. Geographic locations of 6 archaeological sites analysed in this study (red) and four additional Neolithic Anatolian sites relevant for sheep domestication, discussed in the text (black).

Both zooarchaeological data and genomic evidence imply a complex demographic history of domestic sheep. One notable pattern involves the high levels of genetic heterogeneity in domestic sheep. This includes multiple distinct mitochondrial DNA haplogroups found in modern breeds^10^, as well as higher nuclear genetic diversity in sheep compared to that in some other domesticates, such as cattle or dog^11,12^. High diversity would be consistent with scenarios where domestication involved multiple centres and/or a large and heterogeneous wild population. A non-exclusive scenario would be major introgression from wild sheep into domestic flocks, which is supported by zooarchaeological evidence^3^.

Genetic distinctions between Asian and European sheep also imply multiple domestication or wild admixture events. Indeed, present-day sheep cluster in two main groups based on genome-wide polymorphism data: east (Asian and African, including East Mediterranean islands) and west (European)^11,12^. Similarly, Asian and European sheep tend to carry distinct proportions of mitochondrial DNA haplogroups, A and B, respectively^13–15^, a pattern that may have been established already by the 2nd millennium BCE^16,17^.

At the same time, genomic analyses suggest high degrees of allele sharing across domestic sheep breeds. This has been considered evidence for the recent spread of sheep with desired traits across the globe, especially within the last 5 millennia, as part of the secondary products revolution^18,19^. Although the first domesticated sheep were likely used for their meat and possibly their milk^20^, they started to be increasingly exploited for their wool in Bronze Age SW Asia, during the 3rd millennium BCE^21^. Intriguingly, a comparison of DNA retroelements across modern breeds implies an expansion of SW Asian lineages, estimated to date back to the Bronze Age; according to this model, SW Asian sheep with desired traits, such as fine wool, were introduced into local breeds across the globe^22^. A recent ancient DNA study reports evidence consistent with novel breeds being introduced to Bronze Age Europe, coinciding with archaeological evidence for the introduction of wool to this continent^21^. In later periods, export and admixture of selected sheep breeds into local stocks continued^11^. Indeed, the most recent common ancestor of domestic sheep breeds has been inferred to date back only 800 generations ago^11^ - an unexpectedly recent estimate.

We currently lack a solid demographic history model to explain these observations: high diversity, clear genetic structure, and recent coalescence times. What is missing is genetic data on the initial steps of domestication and characterisation of the early domesticated sheep gene pool. Here we present a first attempt to bridge this gap, studying ancient DNA from Neolithic period sheep remains from Anatolia, one of the possible domestication centres. Analysing both mitochondrial DNA (mtDNA) sequences and nuclear polymorphism data, we find support for the notions that the present-day domestic sheep population has multiple origins, and also that the sheep gene pool changed considerably since the Neolithic period.

## Results

We analysed DNA from c.200 archaeological sheep bone and tooth samples from early Holocene Anatolia, originating from six different sites from central and west Anatolia and spanning the Epipaleolithic, Neolithic, and Chalcolithic periods. We obtained and analysed mtDNA sequences from 74 samples, while from four individuals we generated genome-wide ancient DNA data using shotgun sequencing and enrichment capture targeting single nucleotide polymorphisms (SNP). We went on to compare this data with published data sets^11,23^ from present-day wild sheep and domestic sheep breeds (Fig. 1, Supplementary Fig. 1, Supplementary Table 1, Supplementary Table 7).

**Table 1.** High-throughput sequencing summary statistics, AMS C14 ages, molecular sex identifications and mtDNA haplogroups of four ancient sheep. “Genome coverages” were calculated across the full genome length, while the “number of SNPs” indicate those that were covered by at least one read within a set of 40,225 SNPs.

| Sample ID | C14 ages<br>cal. BCE | Genome coverage | Number of SNPs | Molecular sex | mtDNA HPG |
| --- | --- | --- | --- | --- | --- |
| TEP62 | 7031-6687 | 0.273 | 10484 | F | B |
| TEP03 | 7059-6756 | 0.103 | 8223 | F | B |
| TEP83 | 6469-6361 | 0.022 | 3294 | M | A |
| ULU31 | 6227-6071 | 0.022 | 4482 | F | B |

### Mitochondrial DNA data indicates a domestication-related bottleneck around 7500 BCE

To investigate changes in the maternal lineage, we amplified and Sanger sequenced a 144 bp fragment of the mtDNA control region. This region contains diagnostic markers for the five main haplogroups observed in present-day domestic sheep, *i*.*e*. haplogroups A-E^24^, and is short enough to be effectively analysed in ancient samples^23,25^ (Supplementary Table 3). A total of 178 sheep samples were studied, each likely from distinct individuals according to context (Supplementary Materials and Methods). Among these, 74 yielded consistent sequences from at least two independent amplifications. Success rates ranged between 20% to 61% across the six archaeological sites (Supplementary Table 4). We further excluded 4 sequences where diagnostic changes could be confounded by postmortem damage-induced nucleotide transitions (Methods). The sequences thus obtained were analysed to identify haplogroups and haplotypes, and compared across archaeological periods and regions (west and central Anatolia) and with data from present-day sheep breeds (Fig. 2; Supplementary Table 2).

**Fig. 2.**
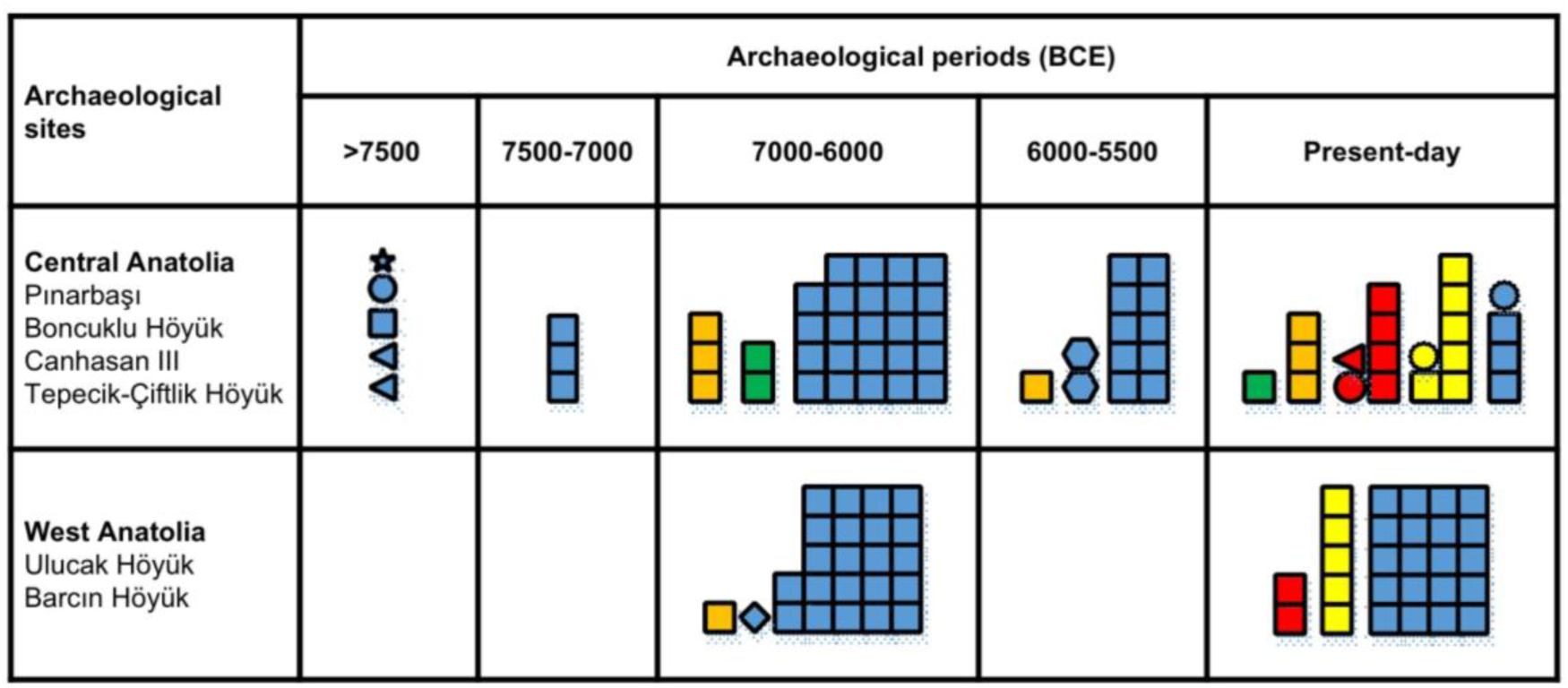
Change in mitochondrial lineages over time. Distribution of different haplogroups (HPG) (indicated by colours) and haplotypes (indicated by geometric shapes) of the studied sheep individuals. Colour coding is as follows: HPG A (red), HPG B (blue), HPG C (yellow), HPG D (green) and HPG E (orange). Note that different periods and haplotypes are not homogeneously represented across different sites from a single region: *e*.*g*. >7500 BCE in central Anatolia is represented by Pinarbasi and Boncuklu Höyük, while 7500-7000 BCE by Canhasan III and Tepecik-Çiftlik Höyük. Meanwhile, the only non-B haplogroup in west Anatolia from 7000-6000 BCE belongs to Ulucak Höyük. See Supplementary Table 2 for full details.

The mtDNA haplogroup data presented in (Fig. 2) reveals a number of interesting patterns. One is the change in haplotype diversity among haplogroup B lineages during the Aceramic Neolithic period, around 7500 BCE. In central Anatolia, haplotype diversity was high (0.9) before 7500 BCE, but totally vanished in the 3 haplotypes we could sample from 7500-6000 BCE. After 6000 BCE diversity rose to 0.3, and then to its present day value of 0.5 (Supplementary Table 5). Notably, haplogroup B appears as the predominant lineage (>90%) in central and west Anatolia, from the Epipaleolithic to the Chalcolithic. Within this group, the specific haplotype that reached 95% (59/62) frequency after domestication (7500-5500 BCE) was already present in the pre-domestication period, but only at 20% (1/5) frequency (Fisher’s exact test p=0.0002). This significant shift in haplotype composition and loss of haplotype diversity in haplogroup B (two-sided permutation test *p*<0.05; Supplementary Fig. 2; Supplementary Table 6) would be consistent with a domestication-related bottleneck during the 8th millennium BCE. Interestingly, we observe that the same haplotype of B that rose in frequency during the Neolithic still appears as the most widespread type today (Fig. 2).

A second pattern involves changes post-7000 BCE, during the Ceramic Neolithic period when farming spreads to west Anatolia and Europe. Compared to pre-7000 BCE, total mtDNA diversity increases in the sample through the appearance of non-B haplogroups (Fig. 2). Although our sample size is yet too small to exclude the presence of non-B haplogroups in central Anatolia pre-7000 BCE, this possible change in haplogroup composition may herald the modest scale introduction of domestic sheep lineages from elsewhere, possibly from another region east of the Fertile Crescent, the south Anatolian coast or the Levant that may have harbored independent domestication events or through ongoing introgression from wild sheep.

Finally, change in haplogroup composition through admixture appears to have continued post-5500 BCE, with significant changes between Neolithic and present-day central Anatolia; this shift happens more subtly in west Anatolia. Overall, analyses of maternal lineages lend support to a domestication event in central Anatolia, as well as major admixture events in the post-Neolithic era sheep populations.

### Anatolian Neolithic sheep show higher genomic affinity to modern European than non-European breeds

We next prepared Illumina high-throughput sequencing libraries from 29 of these ancient sheep samples (Supplementary Table 8). Four Anatolian Neolithic sheep (ANS) individuals’ libraries contained >1% endogenous sheep DNA, with a median of %2. Three were from the central Anatolian site Tepecik-Çiftlik Höyük (TEP03, TEP62, TEP83) and one was from the west Anatolian site Ulucak Höyük (ULU31). The four individuals were AMS C14 dated to the 7th millennium BCE (except for TEP62, for which the age range extended into the 8th millennium), broadly overlapping with the Ceramic Neolithic period in Anatolia (Table 1).

To increase coverage, we enriched the libraries of these four individuals using hybridization capture, targeting 20,000 single nucleotide polymorphisms, and sequenced deeper (Methods). The capture procedure increased the endogenous proportion by 1.5-4x, and resulted in genome coverages ranging between 0.02-0.27x (Table 1). All four libraries exhibited postmortem damage profiles expected for authentic ancient molecules, with >25% C to T transitions at 5’ ends of molecules (Supplementary Fig. 3). After trimming sequencing reads to remove postmortem damage-induced transitions, we called SNPs from these four libraries using the Illumina OvineSNP50 Beadchip variant set^11^, which included 40,225 SNPs mappable to the oviAri3 reference genome. This resulted in a data set containing pseudohaploidised genotypes for 3,294-10,484 autosomal SNPs per individual (Table 1; Methods).

**Figure 3.**
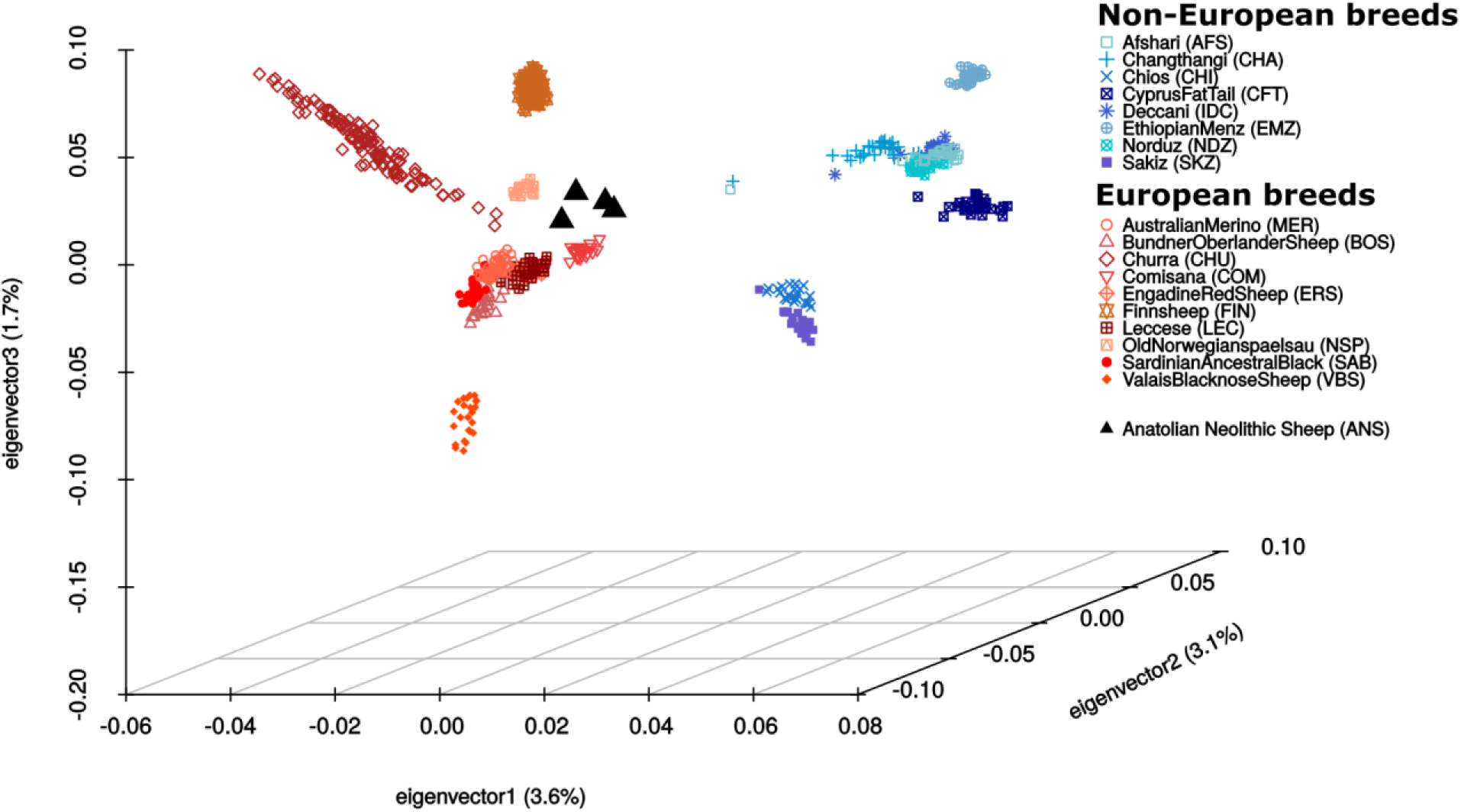
Genomic variation of modern breeds and Anatolian Neolithic sheep. The graph represents the first 3 components of a PCA calculated using genotypes of 18 modern sheep breeds. The four Anatolian Neolithic sheep individuals’ genotypes (triangles) were projected on these 3 components.

We noticed that the average fragment size for the three Tepecik-Çiftlik individuals were >90 bp, uncommonly long for ancient DNA molecules. We used three approaches to investigate possible modern sheep DNA contamination in these libraries: (a) we selected molecules that bear the C->T postmortem damage signature and repeated demographic analyses with only these plausibly authentic molecules; (b) we compared short and long molecules with respect to their postmortem damage signatures; (c) we called genotypes using short and long molecules, performed demographic analyses, and searched for inconsistencies (Methods). None of the results indicated modern DNA contamination (Supplementary Fig. 4-7), leading us to conclude that the long DNA molecules of Tepecik-Çiftlik sheep most probably reflect unusual DNA preservation at this site, consistent with our earlier observations on Neolithic human material from the same site^26^.

**Figure 4.**
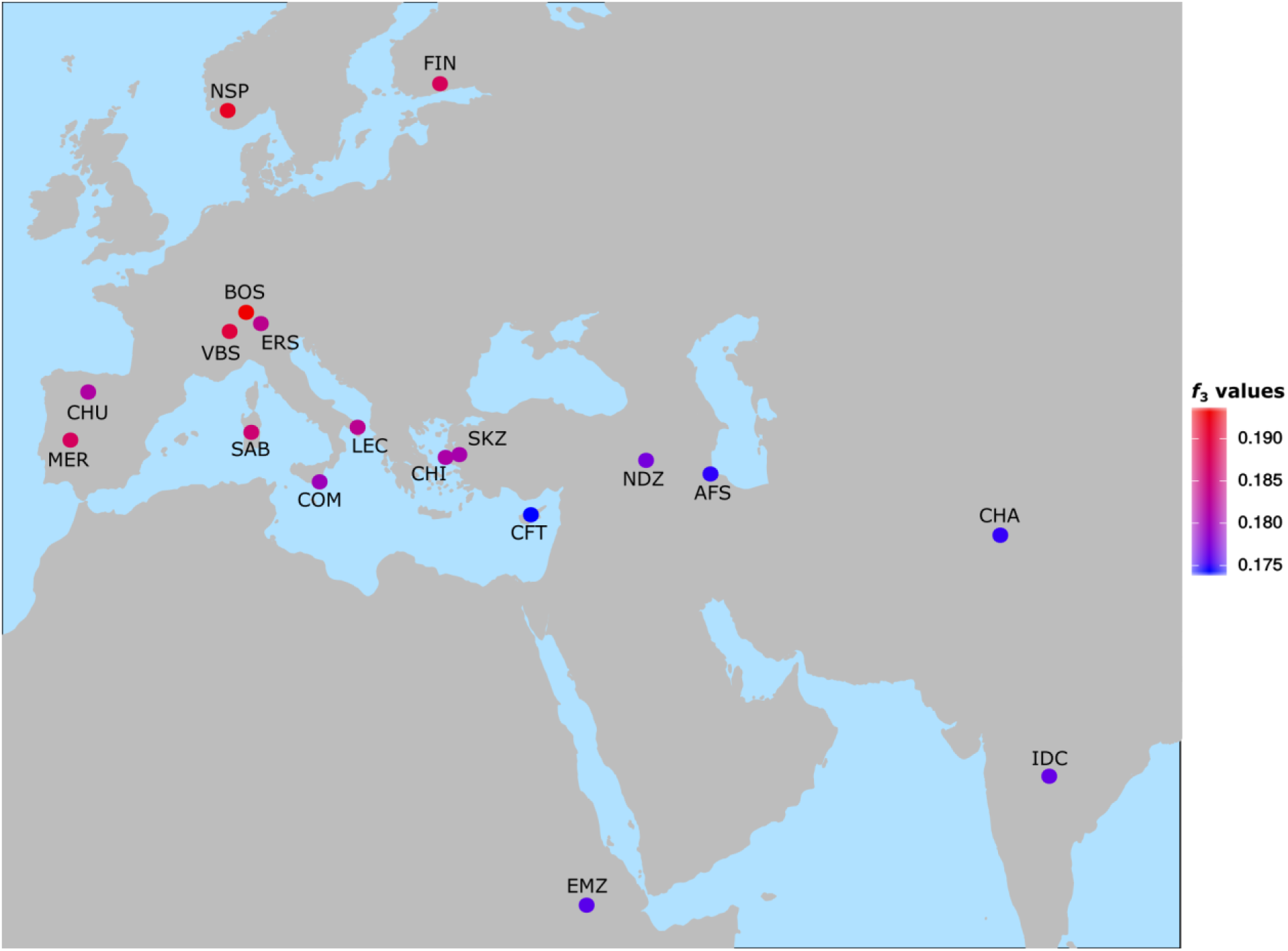
Shared genetic drift between ancient individuals and present-day (modern) populations. Outgroup *f*_*3*_-statistics were calculated as *f*_*3*_(Argali; ANS, modern individual), using the joint allele frequencies of the four ANS individuals. Higher *f*_*3*_ values, in red, indicate higher shared drift.

Next, we sexed the four individuals comparing autosome versus X chromosome coverages, which revealed three females and one male (Table 1, Supplementary Fig. 8). The excess (albeit non-significant) of females is consistent with young male slaughter patterns^9^. We also studied these four individuals’ genotypes at 18 marker SNPs associated with putative domestication-related and positively selected regions, reported by Kijas and colleagues^11^. We found that among 16 loci where we could assign the ancestral state using *Ovis ammon* (Argali) and *Ovis vignei* as outgroups, at 6 loci (38%) the derived allele was carried by at least one Anatolian Neolithic sheep individual, but never among all ANS (Supplementary Table 9). We caution, however, that the present-day linkage disequilibrium between marker alleles and the domestication phenotype-related causal alleles may not necessarily have existed 9,000 years ago. Therefore, this result is not direct evidence that domestication-related derived traits were already present in ANS.

To investigate sheep demographic history with this genome data set, we collected published genomic polymorphism data from ten European breeds (representing north, central and south Europe), and eight non-European breeds from the Middle East, East Mediterranean islands, south Asia, and Africa^11^, generated on Illumina arrays (Supplementary Fig. 1b; Supplementary Table 7). For calculating *f*-statistics we used Argali sheep as outgroup, after confirming that *D*- statistics of the form *D(Goat, Argali; modern*_*1*_; *modern*_*2*_) were all non-significant, suggesting that none of the modern breeds used here had received Argali admixture (1224 tests, multiple testing adjusted *p*>0.05).

We first summarised genome-wide variation through principal component analysis (PCA), calculating the principal components with modern breeds and projecting the four ANS genotypes onto the space described by the first three components (Fig. 3). As observed in earlier work, modern breeds from mainland Europe and of non-European descent (Asian, African, and East Mediterranean) form two distinct clusters in the PCA. Within this world-wide constellation of modern sheep, the ANS attained a relatively central location, although conspicuously closer to the European cluster than to the non-European group.

To confirm this clustering using a formal statistical framework, we calculated *D*-statistics^27^ of the form *D(Argali, ANS*_*1*_; *ANS*_*2*_, *modern)*, where *ANS*_*1*_ and *ANS*_*2*_ denote two different ANS individuals and *modern* denotes any of the 18 present-day (modern) breeds. ANS were consistently genetically closer to each other than to modern breeds (89% of 216 tests, multiple testing adjusted *p*<0.05). Meanwhile, tests of the form *D(Argali, modern; ANS*_*1*_, *ANS*_*2*_) showed that modern breeds did not show higher affinity to any ANS individual over any other ANS individual (216 tests, multiple testing adjusted *p*>0.05). Likewise, *D(Argali, ANS*_*1*_; *ANS*_*2*_, *ANS*_*3*_) showed no significant affinity between any pair of ANS (24 tests, multiple testing adjusted *p*>0.05). These results suggest that Anatolian Neolithic sheep, from different periods and origins, had similar demographic histories.

We further used the outgroup *f*_*3*_ statistic^27^ to measure shared genetic drift between ANS and modern breeds. We calculated *f*_*3*_ in the form of *f*_*3*_*(Argali; modern, ANS)*, where *modern* denotes a modern sheep breed while *ANS* denotes one of the ancient individuals (Methods; Supplementary File 1). The *f*_*3*_ distributions were highly correlated between pairs of ANS individuals (Spearman correlation rho>0.55; Supplementary Fig. 9), indicating that the affinities of ANS to modern breeds were highly alike, irrespective of ANS origin or age. This further supports the notion that ANS were a genetically rather homogeneous population.

The PCA had suggested that ANS may be closer to European modern breeds than to non-European ones. Accordingly, ANS showed significantly higher *f*_*3*_ values with European breeds (median=0.19) than with non-European breeds (median=0.18) (Fig. 4; Mann-Whitney U-test *p*=0.0002), indicating stronger affinity of ANS to present-day European breeds. We further examined this pattern by two approaches. Tests of the form *D(Argali, ANS; European, non*-*European)* revealed that ANS had higher affinity to European breeds (43% of 80 tests were significant, multiple testing adjusted *p*<0.05; Supplementary File 2). This was notable, given that the non-European breeds included east Mediterranean strains that are geographically closest to the ANS individuals’ provenance among all modern breeds analysed. Moreover, in *D*-statistics of the form *D(Argali, ANS; European*_*1*_, *European*_*2*_) ANS showed a trend toward higher allele sharing with central and northern European breeds relative to south European breeds (9% of 90 comparisons had multiple testing adjusted *p*<0.05); in none of these comparisons did we find significantly higher affinity to south European breeds (Supplementary File 2). ADMIXTURE analysis^28^ of modern and ancient breeds likewise indicated similarity between ancestry components in ANS and north and central European breeds (Supplementary Figure 6).

We finally tested modern-ANS relations using *D(Argali, modern*_*1*_; *modern*_*2*_, *ANS)*. Here, in all 306 comparisons performed, modern breeds consistently chose other modern breeds over ANS (multiple testing adjusted *p*<0.05). This result could have a number of explanations, including technical issues and complex demographic histories, which we discuss below.

## Discussion

Our combined analyses of mitochondrial DNA and genome-wide polymorphism data from ancient central and west Anatolian sheep provide novel insights into sheep domestication and later dynamics. First, the abrupt loss of mitochondrial haplotype diversity we observe post-7500 BCE in central Anatolia, and the apparent genetic homogeneity of the four Anatolian Neolithic sheep individuals studied at the genome level, both suggest that we are witnessing signs of domestication in Anatolia. It is also notable that the specific mitochondrial haplotype that becomes common in what are probably the earliest domesticated caprines in the Konya basin is already present in central Anatolia in the Epipalaeolithic, c.6000 years before any morphological or isotope evidence of domestication^29,30^. A central Anatolian domestication scenario would be consistent with archaeological evidence such as early 8th millennium corralling activities documented at the central Anatolian Asikli Höyük^7^, and the mid 8th millennium BCE dramatic shift in sheep and goat diets in the central Anatolian Konya plain^30^.

Second, we observe a trend of increasing haplogroup diversity post-7000 BCE, which could be explained by two non-exclusive models: (a) that another domestication centre existed to the east of central Anatolia (possibly southeast Anatolia and/or the Zagros), the products of which eventually spread westward into central and west Anatolia, and (b) ongoing introgression from wild stocks. Here it is interesting to note that in Anatolian sites dating to the early Chalcolithic (in Çatalhöyük west and in Erbaba in the Lakes district), more than one millennium after the initial decrease in sheep body size, sheep body sizes again rise to wild caprine levels, a pattern that has been interpreted as a sign of wild introgression^3^. Similar patterns have been reported for pigs, cattle and goats, supported by both zooarchaeological analysis^3^ as well as ancient DNA^31,32^. To fully elucidate the history of early sheep domestication, though, we will need to study ancient DNA data from a wider region of southwest Asia.

Third, we find that Anatolian Neolithic sheep show significantly higher affinity to modern-day European breeds than to Asian breeds, including east Mediterranean sheep. This result is also consistent with the mitochondrial haplogroup compositions of ANS and breeds from Europe^13,14,16,33^, where haplogroup B predominates. A possible explanation for this pattern is that ANS were the direct ancestors of modern-day European sheep, and were brought to Europe through the Neolithic migrations of the 7th and 6th millennia^34^. Modern-day Asian sheep, in turn, may have been influenced by non-Anatolian domestic sheep gene pools and/or wild introgression in Asia. Our results further imply that the east Mediterranean and Anatolian sheep gene pools underwent major shifts since the Neolithic, likely through gene flow from the east. Such a turnover would partly echo what has been described for the human gene pool in Anatolia, such that Neolithic Anatolians show higher similarity to modern-day south Europeans than to modern-day Anatolians^26,35^.

Fourth, both *D*-statistics and *f*_*3*_ analyses (Fig. 4) indicate higher affinity of ANS to central and north European sheep than to south European sheep. This observation may suggest that ANS were ancestors of European sheep that followed the Danube (land) route rather than the Mediterranean sea route^36–38^. Alternatively, this pattern could arise due to higher Asian introgression into south than into north European breeds; *e*.*g*. in the post-Neolithic era, Asian alleles could have spread among south European breeds through Mediterranean sea routes. A comparison among present-day breeds supports a scenario of Asian introgression into the Mediterranean: Middle Eastern sheep (AFS, NDZ) are genetically closer to south European than to central or north European breeds [among tests of the form *D(Argali, MiddleEastern; southEuropean, central/northEuropean)*, 25% of 70 comparisons had multiple testing adjusted *p*<0.05]. This raises the possibility that Neolithic and/or post-Neolithic admixture events in the Mediterranean led to the observed higher ANS affinity to central and north European breeds.

An unexpected result here is that in *D*-statistics, modern breeds were consistently closer to other modern breeds than to ANS. Likewise, admixture *f*_*3*_ statistics of the form *f*_*3*_(*southEuropean; ANS, MiddleEastern*) did not yield significantly negative results (*p*>0.05), which would have been expected if South European breeds were a product of simple admixture between ANS and Middle Eastern breeds (Supplementary File 1). One possible explanation could be technical: while the modern data is based on arrays, the ANS data is based on shotgun sequencing and also capture; either technology may be biased with respect to alleles genotyped (Methods). Yet another, biological explanation could be post-Neolithic admixture events that universally influenced all sheep breeds, eclipsing earlier trends. For instance, a west Asian sheep lineage bred for its fine wool may have spread during and after the Chalcolithic and dramatically influenced the global sheep gene pool, which would be consistent with the aforementioned high degrees of haplotype sharing^11^ or retrovirus genotype sharing^22^ observed among modern-day breeds, as well as recent ancient DNA work implicating post-Neolithic gene flow from eastern sources altering the west Eurasian sheep gene pools^21^. Our results suggest that although central Anatolian wild sheep were probably locally domesticated and eventually gave rise to Europe’s first domestic sheep, the present-day domestic sheep gene pool was strongly remoulded by subsequent admixture events of Asian origin.

## Material and Methods

### Sample collection

Ancient sheep bone and tooth samples were obtained from 6 archaeological sites: Pinarbasi, Boncuklu Höyük, Tepecik-Çiftlik Höyük, and Canhasan III in central Anatolia, and Ulucak Höyük and Barcın Höyük in west Anatolia (Fig. 1). Brief information about the sites are provided in Supplementary Material and Methods.

### AMS radiocarbon dating

Five samples were AMS C14 dated at the TÜBİTAK-MAM (Gebze, Turkey) and one sample at Beta Analytic Inc. (London, UK). Radiocarbon ages were calibrated using the INTCAL13 database. The 2 sigma calibrated age estimates were as follows: TEP3_depo: 7059-6756 BCE (TÜBİTAK-694), TEP58: 6645-6505 BCE (Beta-373271), TEP62: 7031-6687 BCE (TÜBİTAK-695), TEP83: 6469-6361 BCE (TÜBİTAK-696), and ULU31: 6227-6071 BCE (TÜBİTAK-697), respectively (Supplementary Table 1). The remaining samples were dated by the excavation directors based on their archaeological context.

### Ancient DNA extraction

Ancient DNA extraction was performed in a dedicated aDNA laboratory at METU, following the protocol described in Dabney et al.^39^ (Supplementary Material and Methods). DNA was extracted twice from each sample at different times.

### mtDNA sequencing and haplogroup assignment

The 144 bp long fragment of sheep mtDNA corresponding to the positions 15391-15534 on the reference AF010406 sequence was sequenced from 76 ancient samples using published primer pairs^25^. Samples were assigned to mtDNA haplogroups (A to E) according to the identity of nucleotides on haplogroup-determining positions with respect to the reference AF010406 sequence (Supplementary Table 3). Following shotgun sequencing, we determined that one individual’s assignment was inconsistent between mtDNA and Illumina sequencing data, which we determined to be caused by postmortem damage at mtDNA sequences. We consequently corrected one haplogroup assignment based on Illumina sequencing, and we further removed three sequences (all haplogroup A) where assignment could be confounded by postmortem damage.

### Mitochondrial genetic diversity

Genetic diversity measures such as haplogroup and haplotype diversity were calculated using *DnaSP* (v.6)^40^ and their significance were determined by random permutation tests (Supplementary Material and Methods).

### Whole genome libraries and prescreening

We prepared 36 double-stranded Illumina sequencing libraries following Meyer and Kircher^41^ and sequenced these on Illumina HiSeq platforms at low coverage (median c.13 million reads per library) (Supplementary Table 8). Libraries from four individuals (TEP3, TEP62, TEP83, ULU31) contained >1% endogenous sheep DNA, while other libraries had negligible proportions.

### Hybridization capture

To increase genome coverage, the chosen four libraries were used for hybridization capture with custom designed 80K probes targeting 20K SNPs. Briefly, the SNPs were chosen from the Illumina OvineSNP50 Beadchip variant set, giving priority to transversions and also including mitochondrial markers and SNPs associated with putatively positively selected regions^11^ (Supplementary Material and Methods). The biotinylated RNA capture probes were produced by Arbor Biosciences Inc. and capture experiments were implemented following the manufacturer’s instructions.

### Data preprocessing

We combined BAM files from shotgun and capture libraries from the same individual, removed the residual adapter sequences in .*fastq* files and merged paired-end sequencing reads using *MergeReadsFastQ_cc*.*py*^42^. We mapped the merged reads to the sheep reference genome (Oar_v3.1) using *BWA aln* (v. 0.7.12)^43^, merged all libraries from the same individual using *SAMtools merge*^44^, removed the PCR duplicates using *FilterUniqueSAMCons*.*py*^42^, removed reads shorter than 35 base pairs and/or with >10% mismatches to the sheep reference genome. Ancient individuals’ *BAM* files were trimmed from both ends by 10 bp using *trimBAM* command of *bamUtil* software^45^ to avoid postmortem damage at read ends being interpreted as true variants. We used the Illumina OvineSNP50 Beadchip SNP panel for genotype calling, with 40,225 SNPs in this list that could be mapped to the oviAri3 reference sequence. We ran the *SAMtools* (v. 1.3)^46^ *mpileup* program on *BAM* files and pseudohaploidised the data by randomly choosing a single read to represent the genotype^47^ (Supplementary Material and Methods).

### Authentication of ancient sequences

We used *PMDtools*^48^ to measure postmortem damage patterns at 5’ and 3’ ends of the reads, and generated postmortem damage profile graphs using the *PMDtools* ‘*--deamination*’ parameter. Observing read lengths longer than usual (>90 bp) in 3 Tepecik-Çiftlik libraries, we used multiple approaches to rule out modern sheep DNA contamination. (1) We selected 27-53% (median 41%) DNA molecules bearing the postmortem damage signature (i.e. were most likely authentic) using PMDtools^48^ with the *‘--threshold 3’* parameter. We then compared the PMD-bearing reads with the unfiltered read set with respect to read lengths (Supplementary Figure 4); we observed that these molecules were not shorter than the rest, which would have been expected if the long reads represented modern DNA contamination. (2) We repeated PCA and outgroup *f*_*3*_ calculations of ANS using these PMD-bearing reads only (Supplementary Figure 5); we found that using this restricted read set yields the same fundamental observations as using all reads. (3a) From each individual’s BAM files we selected short (<70 bp) and long (>100 bp) reads. We compared 5’ C->T damage profiles between these short vs. long read sets, which did not reveal any systematic difference (Supplementary Figure 6). (3b) We called SNPs from these short vs. long read sets: 1616 vs. 7661 for TEP03, 2817 vs. 9318 for TEP62, 544 vs. 2009 for TEP83, and 3547 vs. 500 for ULU31. Using these datasets, we calculated outgroup *f*_*3*_ of the form *f*_*3*_(*Argali; ANS*_*i*_, *modern*), where *ANS*_*i*_ represents one of the ANS individual’s genotype based on either short or long reads, and *modern* represents a modern sheep breed’s genotype. We then calculated the Pearson correlation between outgroup *f*_*3*_ values based on short vs. long reads for each individual. The correlation coefficients were all positive, although significant for only TEP62 and ULU31 (Supplementary Figure 7). In addition, in the main text, we show that all four ANS individuals, irrespective of provenance or read length, displayed similar population genetic affinities, which further supports the notion that the long molecules detected in Tepecik-Çiftlik are authentic.

### Molecular sex determination

We determined the molecular sex of samples by comparing the relative mapping frequency of autosomes to the X chromosome using regression analysis (Supplementary Material and Methods).

### Modern sheep genotypes

From the 74 worldwide breeds included the Kijas et al.^11^ SNP chip dataset, we chose a subset that would be representative and relevant to our study, and also exclude breeds known to have undergone strong bottlenecks (e.g. Soay) or recent admixture (e.g. Creole). We thus selected 10 European (north, middle and south) and 8 non-European modern breeds, the latter from the Middle East, south Asia, Africa, and the east Aegean Sea (Supplementary Table 7).

### Principal component analysis

We merged four ancient individuals with the chosen 18 modern breeds using *PLINK*^49^. We conducted principal component analysis (PCA) using the *smartpca* command of *EIGENSOFT* (v. 7.2.0) software^50^. Components of eighteen modern populations and four Mouflon individuals from SheepHapMap project dataset were first calculated, and the four ANS individuals were projected onto the first three components (Fig. 3). Visualization of the PCA was done in the *R* (v. 3.5) environment with the *scatterplot3d* package^51^.

### *f*_*3*_-statistics

We performed outgroup-*f*_*3*_ and admixture-*f*_*3*_ statistics using the *qp3Pop* program of the *AdmixTools* (v. 5.1) software27. We performed outgroup-*f*_*3*_ in the form *f*_*3*_*(Argali; ANS, modern)*, where *Argali* represents the outgroup (for which we randomly chose one Argali sheep individual), *ANS* represents the genotype of an Anatolian Neolithic sheep individual or all four Anatolian Neolithic sheep combined, and *modern* represents the genotype one of the eighteen modern breeds. In order to calculate admixture *f*_*3*_-statistics, we repeated the same procedure but using the *Admixtools* (v. 5.1) software^27^ with the *‘inbreed: YES*’ parameter, and calculating *f*_*3*_*(modern1; ANS, modern2)*, where *ANS* represents the genotype of all four Anatolian Neolithic sheep combined, and *modern1* and *modern2* represent the genotypes of modern south European and modern Asian breeds, respectively.

### *D*-statistics

We conducted *D*-statistics using the *qpDstat* program of *AdmixTools* (v. 5.1) software^27^. For this, we constructed subsets of the data set used for principal component analysis. As in the outgroup-*f*_*3*_ analysis, we used the same Argali sheep individual as outgroup. To control for the false positive rate, we performed multiple testing correction using Benjamini and Hochberg method^52^ in the *R* (v. 3.5) environment separately for each set of comparisons.

### ADMIXTURE analysis

We conducted clustering analysis using *ADMIXTURE* (v.1.3) software^28^. We pruned the data set to remove SNPs in linkage disequilibrium using *PLINK*^49^, and performed 10 trials for all *K*’s between 2 and 12. We used the *Pong* software^53^ to visualize *ADMIXTURE* results (Supplementary Material and Methods).

## Data availability

All .*fastq* files were submitted to the European Nucleotide Archive (ENA) with reference number PRJEB36540.

## Code availability

The code for probe design is available at https://github.com/dkoptekin/bait-design.

## Acknowledgements

We are grateful to Torsten Günther, Pedro Morell Miranda and other members of the Günther Lab and Daniel Bradley for helpful suggestions and/or comments. This work was supported by TÜBİTAK 1001 (Project No: 114Z356) and ERC Consolidator grant “NEOGENE” (Project No 772390).

## Author contributions

I. Togan, M.S., F.Ö., A.G., designed and supervised the study; D.B., Y.G.Ç., E.B., Ö.Ç., F.G., L.M., R.Ö., E.P., equally contributed archaeological samples, archaeological information, discussion and writing up of results; C.Y.G, M.Ş., J.H., equally contributed archaeological samples and archaeological information; O.Ö., E.Yüncü., N.D.D., A.A., S.Y., V.K.L., M.K., F.A., F.Ö., performed the wet lab experiments; E.Yurtman., O.Ö., D.Ko., D.Ka., G.A., performed data analysis with contributions from M.Ö., K.B.V., G.M.K., A.G., S.C.A., R.Y., E.S., C.C.B., A.G., M.S., İ.T.; and M.S., İ.T., E.Yurtman., O.Ö., F.Ö., D.Ko., wrote the manuscript with input from all authors.

