## Supplemental Information for "Archaeogenetic analysis of Neolithic sheep from Anatolia suggests a complex demographic history since domestication"

### Supplementary Tables

**Supplementary Table 1.** Genetic and archaeological information on sheep individuals used in mitochondrial DNA analyses.

The table shows names of the archaeological sites for samples studied, lab and excavation site identification numbers, approximate age by context, number of times independent sequences were obtained, and the mitochondrial haplogroups (HPG) assigned to each sample. Genome data is available for samples marked with asterisks (*) in the second column (see Supplementary Table 6). HPG: Haplogroup. TEP3_depo, TEP58, TEP62 and ULU31 was directly dated in this study (shown in bold). Ages marked with δ are conventional radiocarbon ages before present (BP).

| Archaeological Site | Lab ID | Excavation IDs | Date | Number of times sequenced | HPG |
|---|---|---|---|---|---|
| Pınarbaşı | PB2 | ABU 193 | 9600-7800 cal BC | 3 | B |
| Pınarbaşı | PB10 | BBH 10208 | 6500-6000 cal BC | 2 | D |
| Pınarbaşı | PB12 | BBH 10025 | 6500-6000 cal BC | 2 | B |
| Pınarbaşı | PB25 | BHL 8688 | 14000-11000 cal BC | 2 | B |
| Pınarbaşı | PB28 | BIE 8194 | 14000-11000 cal BC | 2 | B |
| Pınarbaşı | PB33 | BIH 4094 | 14000-11000 cal BC | 2 | B |
| Pınarbaşı | PB6 | BBH 188 | 6500-6000 cal BC | 2 | B |
| Pınarbaşı | PB8 | BBH 10207 | 6500-6000 cal BC | 2 | B |
| Boncuklu Höyük | BK3 | NAE 2945 | 8400-7800 cal BC | 2 | B |
| Canhasan III Höyüğü | CH1 | 49F BONE ID 142 | 7400-7100 cal BC | 2 | B |
| Canhasan III Höyüğü | CH2 | 49L BONE ID 137 | 7400-7100 cal BC | 2 | B |
| Canhasan III Höyüğü | CH3 | 49T BONE ID 087 | 7400-7100 cal BC | 2 | B |
| Tepecik-Çiftlik Höyük | TEP1 | 1/15J/52C | 6000-5800 BCE | 3 | B |
| Tepecik-Çiftlik Höyük | TEP10 | 10/15K/101C | 6000-5800 BCE | 2 | E |
| Tepecik-Çiftlik Höyük | TEP17 | 17/15K/111C | 6000-5800 BCE | 2 | B |
| Tepecik-Çiftlik Höyük | TEP20 | 20/15K/109C | 6000-5800 BCE | 2 | B |
| Tepecik-Çiftlik Höyük | TEP3_depo* | 780/16K | **7059-6756 cal BCE (C14) (7994+/-41 BP)**$^{\delta}$ | 2 | B |
| Tepecik-Çiftlik Höyük | TEP4 | 4/15K/100C | 6000-5800 BCE | 2 | B |
| Tepecik-Çiftlik Höyük | TEP41 | 41/15K/122C | 6000-5800 BCE | 3 | B |
| Tepecik-Çiftlik Höyük | TEP43 | 43/15J/87C | 6000-5800 BCE | 3 | B |
| Tepecik-Çiftlik Höyük | TEP45 | 45/15K/119C | 6000-5800 BCE | 2 | B |
| Tepecik-Çiftlik Höyük | TEP46 | 46/15K/119C | 6000-5800 BCE | 2 | B |
| Tepecik-Çiftlik Höyük | TEP49 | 49/15K/65C | 6000-5800 BCE | 2 | B |
| Tepecik-Çiftlik Höyük | TEP4_depo | 779/16K | 7100-6900 BCE | 3 | B |
| Tepecik-Çiftlik Höyük | TEP50 | 50/15K/89C | 6000-5800 BCE | 2 | B |

| | | | | | |
|---|---|---|---|---|---|
| Tepecik-Çiftlik Höyük | TEP52 | 52/16K/247C | 6700-6600 BCE | 2 | B |
| Tepecik-Çiftlik Höyük | TEP53 | 53/16K/245C | 6700-6600 BCE | 3 | B |
| Tepecik-Çiftlik Höyük | TEP54 | 54/16K/253C | 6700-6600 BCE | 3 | B |
| Tepecik-Çiftlik Höyük | TEP55 | 55/16K/263C | 6700-6600 BCE | 3 | E |
| Tepecik-Çiftlik Höyük | TEP56 | 56/16K/265C | 6700-6600 BCE | 2 | E |
| Tepecik-Çiftlik Höyük | TEP57 | 57/16K/265C | 6700-6000 BCE | 3 | B |
| Tepecik-Çiftlik Höyük | TEP58 | 58/16K/265C | **6645-6505 cal BCE (C14) (7760+/-30 BP)**[δ] | 2 | B |
| Tepecik-Çiftlik Höyük | TEP60 | 60/16K/267C | 6700-6000 BCE | 3 | B |
| Tepecik-Çiftlik Höyük | TEP61 | 61/16K/262C | 6800-6700 BCE | 3 | B |
| Tepecik-Çiftlik Höyük | TEP62* | 62/16K/262C | **7031-6687 cal BCE (C14) (7934+/-37 BP)**[δ] | 2 | B |
| Tepecik-Çiftlik Höyük | TEP7 | 7/15K/100C | 6000-5800 BCE | 3 | B |
| Tepecik-Çiftlik Höyük | TEP7_depo | 783/16L | 6500-6400 BCE | 3 | D |
| Tepecik-Çiftlik Höyük | TEP8 | 8/15K/100C | 6000-5800 BCE | 2 | B |
| Tepecik-Çiftlik Höyük | TEP8_depo | 783/16L | 6500-6400 BCE | 3 | B |
| Tepecik-Çiftlik Höyük | TEP12_depo | -/16K | 6800-6600 BCE | 2 | B |
| Tepecik-Çiftlik Höyük | TEP16_depo | -/16K | 6800-6600 BCE | 2 | E |
| Tepecik-Çiftlik Höyük | TEP48_2014 | 1345/17K/280C | 6800-6600 BCE | 2 | B |
| Tepecik-Çiftlik Höyük | TEP57_2014 | 1353/16K/173C | 6700-6600 BCE | 2 | B |
| Tepecik-Çiftlik Höyük | TEP60_2014 | 1356/16K/250C | 6700-6600 BCE | 2 | B |
| Tepecik-Çiftlik Höyük | TP68_2014 | 1362/16K/276C | 6900-6800 BCE | 2 | B |
| Tepecik-Çiftlik Höyük | TEP72_2014 | 1365/16K/267C | 6900-6800 BCE | 3 | B |
| Tepecik-Çiftlik Höyük | TEP82_2014 | 1372/17K/229C | 6700-6600 BCE | 2 | B |
| Tepecik-Çiftlik Höyük | TEP87_2014 | 1377/17K/229C | 6800-6700 BCE | 2 | B |
| Tepecik-Çiftlik Höyük | TEP93_2014 | 1383/17K/5-6 210C | 6800-6700 BCE | 2 | B |
| Tepecik-Çiftlik Höyük | TEP94_2014 | 1384/17K/6- 226C | 6800-6700 BCE | 2 | B |
| Ulucak Höyük | ULU12 | Ulucak 8, VI, Astv | 7000-6500 BCE | 3 | B |
| Ulucak Höyük | ULU27 | Ulucak 5, VI, MD | 7000-6500 BCE | 2 | E |
| Ulucak Höyük | ULU31* | L12C, 139, KNU | **6227-6071 cal BCE (C14) (7291+/-38 BP)**[δ] | 3 | B |
| Ulucak Höyük | ULU32 | L12D, 168, 215.13, 215.07, LHU | 6500-6000 BCE | 2 | B |
| Ulucak Höyük | ULU34 | L12, ctd, Birim 141, 215.11, 215.08, KOG | 6500-6000 BCE | 3 | B |
| Ulucak Höyük | ULU36 | L12D, 163, 215.16, 215.07, LEB | 6500-6000 BCE | 2 | B |
| Ulucak Höyük | ULU4 | Ulucak 7, VI | 7000-6500 BCE | 3 | B |

| | | | | | |
|---|---|---|---|---|---|
| Ulucak Höyük | ULU6 | Ulucak 16, VI | 7000-6500 BCE | 3 | B |
| Barcın Höyük | BH12 | 26485 | 6300-6200 BCE | 3 | B |
| Barcın Höyük | BH14 | 30790 | 6300-6200 BCE | 3 | B |
| Barcın Höyük | BH15 | 30418 | 6500-6400 BCE | 2 | B |
| Barcın Höyük | BH16 | 26474 | 6300-6200 BCE | 3 | B |
| Barcın Höyük | BH17 | 37302 | 6300-6200 BCE | 3 | B |
| Barcın Höyük | BH19 | 37301 | 6500-6400 BCE | 2 | B |
| Barcın Höyük | BH2 | 26477 | 6500-6400 BCE | 2 | B |
| Barcın Höyük | BH20 | 37303 | 6500-6300 BCE | 2 | B |
| Barcın Höyük | BH24 | 37339 | 6300-6200 BCE | 2 | B |
| Barcın Höyük | BH26 | 26488 | 6300-6100 BCE | 2 | B |
| Barcın Höyük | BH27 | 31319 | 6500-6300 BCE | 3 | B |
| Barcın Höyük | BH3 | 26484 | 6300-6100 BCE | 3 | B |
| Barcın Höyük | BH32 | 37084 | 6300-6200 BCE | 2 | B |
| Barcın Höyük | BH41 | 37078 | 6300-6200 BCE | 3 | B |
| Barcın Höyük | BH6 | 31083 | 6300-6200 BCE | 2 | B |
| Barcın Höyük | BH7 | 31183 | 6300-6200 BCE | 2 | B |

**Supplementary Table 2.** Mitochondrial haplogroups (HPG) of three present-day sheep breeds from Anatolia.

The table shows haplogroup frequencies among present-day sheep breeds from Anatolia, published by Demirci et al.[1]. To compare with ancient mitochondrial haplogroup frequencies generated in the current study, we restricted the modern data to those individuals whose 144 bp sequence (the fragment used in the ancient analyses) was available. The modern individuals represented in Figure 2 are thus a subset of the individuals in this table. Sakız originates from West Anatolia, Kıvırcık from Northwest Anatolia, and Akkaraman from Central Anatolia.

| **Breeds** | **HPG A (%)** | **HPG B (%)** | **HPG C (%)** | **HPG D (%)** | **HPG E (%)** |
|---|---|---|---|---|---|
| Sakız (*n*=49) | 4 | 88 | 8 | 0 | 0 |
| Kıvırcık (*n*=45) | 0 | 96 | 4 | 0 | 0 |
| Akkaraman (*n*=50) | 26 | 52 | 14 | 2 | 6 |

**Supplementary Table 3.** Mitochondrial DNA control region marker variants.

mtDNA nucleotide positions used for haplogroup assignment, based on coordinates of the *Ovis aries* mitogenome AF010406 (NCBI GenBank). Table adapted from Demirci et al.[1].

| Haplogroup | Reference sequence | Positions and identities of bases used for haplogroup assignment | | | | |
|---|---|---|---|---|---|---|
| | | 15459 | 15476 | 15484 | 15509 | 15512 |
| HPG A | HM236174 | T | T | A | A | T |
| HPG B | HM236176 | C | T | G | A | T |
| HPG C | HM236178 | C | T | G | G | T |
| HPG D | HM236180 | C | T | G | A | C |
| HPG E | HM236182 | C | C | G | G | T |

**Supplementary Table 4.** Mitochondrial DNA amplification and sequencing success rates.

Success refers to aDNA fragments being amplified and sequenced at least twice, independently.

| Archaeological site | Number of samples studied | Number of successful samples | Success rate |
|---|---|---|---|
| Ulucak Höyük | 40 | 8 | 29% |
| Barcın Höyük | 44 | 17 | 39% |
| Canhasan III | 5 | 3 | 60% |
| Tepecik-Çiftlik Höyük | 67 | 41 | 61% |
| Boncuklu Höyük | 5 | 1 | 20% |
| Pınarbaşı | 17 | 8 | 47% |
| TOTAL | 178 | 78 | 41% |

**Supplementary Table 5.** Mitochondrial genetic diversity of ancient sheep across time.

Haplogroup diversity (*H*) was calculated using the Shannon diversity index and including all samples within a group. The haplotype diversity (*hd*) and standard deviation estimates were calculated using samples belonging to haplogroup B only, with the *DnaSP* software[2]. Empty cells have no data available. The significance of *H* and *hd* differences among periods was calculated using permutation tests. The significance of *hd* differences among periods are shown in Supplementary Table 6, while none of the *H* differences were significant among periods.

| **Archaeological sites** | **Statistic** | **Period (BCE)** | | | | |
|---|---|---|---|---|---|---|
| | | **>7500** | **(7500 - 7000)** | **(7000 - 6000)** | **(6000 - 5500)** | **Present-day** |
| **Central Anatolia**<br>Pınarbaşı<br>Boncuklu Höyük<br>Canhasan III<br>Tepecik-Çiftlik Höyük | ***H*** | 0 | 0 | 0.58 | 0.27 | 1.21 |
| | ***hd*** | 0.9 ± 0.16 | 0 | 0 | 0.3 ± 0.15 | 0.5 ± 0.27 |
| **West Anatolia**<br>Ulucak Höyük<br>Barcın Höyük | ***H*** | - | - | 0.17 | - | 0.54 |
| | ***hd*** | - | - | 0.09 ± 0.08 | - | 0 |

**Supplementary Table 6** Haplotype diversity (*hd*) differences between different periods.

We calculated differences in *hd* levels between different periods and regions. We then estimated the significance of these differences by performing a permutation (randomization) test of the null hypothesis that diversity differences between two periods were solely by chance (see Supplementary Materials and Methods). The table shows the differences and the two-sided permutation test *p*-values in parentheses. CA: Central Anatolia; WA: West Anatolia; following the legend of Supplementary Table 5.

| | **CA<br>(7500 - 7000) BCE** | **CA<br>(7000 - 6000) BCE** | **WA<br>(7000 - 6000 ) BCE** | **CA<br>(6000 - 5500) BCE** |
|---|---|---|---|---|
| **CA<br>>7500 BCE** | 0.90 ($p$=0.07) | 0.90 ($p$<0.01) | 0.81 ($p$<0.01) | 0.60 ($p$<0.01) |
| **CA<br>(7500 – 7000) BCE** | - | 0 (n.s.) | 0.09 (n.s.) | -0.30 (n.s.) |
| **CA<br>(7000 - 6000) BCE** | - | - | - | -0.30 (n.s.) |
| **WA<br>(7000 – 6000) BCE** | - | - | - | -0.22 ($p$=0.07) |

**Supplementary Table 7.** Modern sheep breeds used in this study in genomic analyses. Breed names, abbreviations and geographic origins of present-day breeds, from Kijas et al.[3].

| **Abbreviation** | **Breed Name** | **Origin** |
|---|---|---|
| AFS | Afshari | SW Asia |
| BOS | Bundner Oberlander Sheep | Central Europeæ |
| CFT | Cyprus Fat Tail | SW Asia |
| CHA | Changthangi | S Asia |
| CHI | Chios | SW Europe |
| CHU | Churra | SW Europe |
| COM | Comisana | SW Europe |
| EMZ | Ethiopian Menz | Africa |
| ERS | Engadine Red Sheep | Central Europe |
| FIN | Finn sheep | Northern Europe |
| IDC | Deccani | S Asia |
| LEC | Leccese | SW Europe |
| MER | Australian Merino | SW Europe |
| NDZ | Norduz | SW Asia |
| NSO/NSP | Old Norwegian Spælsau | Northern Europe |
| SAB | Sardinian Ancestral Black | SW Europe |
| SKZ | Sakiz | SW Asia |
| VBS | Valais Blacknose Sheep | Central Europe |

**Supplementary Table 8.** Sequencing and mapping statistics for ancient sheep DNA prescreening and capture libraries.

The table presents sequencing and mapping statistics for ancient sheep Illumina libraries sequenced on the Illumina HiSeq platform. The samples include material from 3 sites: Ulucak Höyük (uhs), Tepecik-Çiftlik Höyük (tps) and Barcın Höyük (bhs). The reported sheep proportions were calculated based on the proportion of all reads mapping to the sheep genome, before filtering for mismatches and removing duplicates. Sequencing IDs marked by asterisks ($n$=4) refer to SNP capture libraries.

| Sequencing ID | Sample ID | Excavation ID | Number of reads | Reads mapping to sheep | Sheep proportion | Mean read length | Genome coverage | mtDNA coverage |
|---|---|---|---|---|---|---|---|---|
| bhs003 | BH3 | 26484 | 14285758 | 778 | 0.00023 | 81.4 | 0.000022 | 0.0037 |
| bhs004 | BH4 | 26487 | 12059036 | 979 | 0.00087 | 75.8 | 0.000025 | 0.0070 |
| bhs007 | BH7 | 31183 | 10764539 | 10279 | 0.00159 | 58.5 | 0.000202 | 0.0050 |
| bhs015 | BH15 | 30418 | 24854171 | 3753 | 0.00037 | 69.0 | 0.000089 | 0.0126 |
| tps001 | TEP03 | 780/16K | 9009099 | 160032 | 0.02118 | 105.1 | 0.006185 | 0.3423 |
| tpsc001* | TEP03 | 780/16K | 94338393 | 2096292 | 0.09852 | 126.4 | 0.097290 | 1.8154 |
| tps002 | TEP02 | 779/16K | 8954211 | 51829 | 0.00745 | 51.8 | 0.000814 | 0.0939 |
| tps005 | TEP05 | 780/16K | 12388414 | 2386 | 0.00034 | 74.4 | 0.059620 | 0.0139 |
| tps009 | TEP09 | NA | 6835967 | 553 | 0.00021 | 79.4 | 0.000015 | 0 |
| tps010 | TEP10 | NA | 5616596 | 10641 | 0.00219 | 108.6 | 0.000400 | 0.0174 |
| tps017 | TEP17 | 17/15K/111C | 13077456 | 91095 | 0.00817 | 96.6 | 0.003157 | 0.8109 |

| | | | | | | | | |
|---|---|---|---|---|---|---|---|---|
| tps020 | TEP20 | 20/15K/109C | 18068470 | 10319 | 0.00068 | 81.9 | 0.000282 | 0.0288 |
| tps046 | TEP46 | 46/15K/119C | 11910234 | 936 | 0.00018 | 85.7 | 0.000029 | 0 |
| tps060 | TEP60 | 60/16K/267C | 17667489 | 6082 | 0.00065 | 101.4 | 0.000217 | 0.0969 |
| tps062 | TEP62 | 62/16K/262C | 15288906 | 1492645 | 0.11534 | 102.1 | 0.056297 | 5.5566 |
| tpsc062* | TEP62 | 62/16K/262C | 74121812 | 5356405 | 0.21889 | 112.3 | 0.221251 | 10.4074 |
| tps083 | TEP83 | 1373/17K/5-6 212C | 15562236 | 147274 | 0.01034 | 84.1 | 0.004174 | 1.7839 |
| tpsc083* | TEP83 | 1373/17K/5-6 212C | 71880314 | 558736 | 0.04121 | 101.5 | 0.019220 | 5.0523 |
| uhs004 | ULU04 | Ulucak 7, VI | 11590200 | 991 | 0.00021 | 70.1 | 0.000024 | 0 |
| uhs006 | ULU06 | Ulucak 16, VI | 10440876 | 1404 | 0.00033 | 85.5 | 0.000042 | 0 |
| uhs009 | ULU09 | NA | 26735725 | 5004 | 0.00078 | 74.9 | 0.000131 | 0.0131 |
| uhs012 | ULU12 | Ulucak 8, VI, Astv | 12464722 | 98615 | 0.00910 | 60.4 | 0.002104 | 0.1355 |
| uhs016 | ULU16 | Ulucak 2, VI, Astr | 15986774 | 2142 | 0.00096 | 64.6 | 0.000047 | 0.0064 |
| uhs023 | ULU23 | Ulucak 19, IV, HM | 8440745 | 184134 | 0.02536 | 60.6 | 0.003868 | 0.1295 |

| | | | | | | | | |
|---|---|---|---|---|---|---|---|---|
| uhs026 | ULU26 | Ulucak 3, VI, M3 | 18816066 | 22817 | 0.00259 | 58.9 | 0.000453 | 0.0770 |
| uhsc026* | ULU26 | Ulucak 3, VI, M3 | 89046667 | 123186 | 0.00376 | 72.2 | 0.003035 | 0.3231 |
| uhs027 | ULU27 | Ulucak 5, VI, MD | 8792111 | 5457 | 0.00104 | 60.8 | 0.000116 | 0.0052 |
| uhs030 | ULU30 | L126, 125, 215.36, 215.15, KKL | 13197914 | 826 | 0.00036 | 77.9 | 0.000022 | 0.0091 |
| uhs031 | ULU31 | L12C, 139, KNU | 15367439 | 325243 | 0.02591 | 53.0 | 0.006013 | 0.0916 |
| uhsc031* | ULU31 | L12C, 139, KNU | 75840411 | 870949 | 0.04813 | 58.6 | 0.017590 | 0.1711 |
| uhs032 | ULU32 | L12D, 168, 215.13, 215.07, LHU | 14645301 | 1090 | 0.00080 | 80.5 | 0.000029 | 0.0192 |
| uhs034 | ULU34 | L12, ctd, Birim 141, 215.11, 215.08, KOG | 14256598 | 23868 | 0.00295 | 54.5 | 0.000422 | 0.1338 |
| uhs036 | ULU36 | L12D, 163, 215.16, 215.07, LEB | 13706310 | 4750 | 0.00107 | 61.5 | 0.000094 | 0.0032 |
| uhs037 | ULU37 | L12, ctd, 94, 215.02, 214.50, LHP | 13701446 | 3496 | 0.00096 | 58.3 | 0.000067 | 0.0029 |

**Supplementary Table 9.** Phenotype-related SNPs in ancient individuals.

Reference allele frequencies across *n*=749 modern domestic, *n*=6 Argali and *n*=4 Vignei sheep, as well as genotypes of four Anatolian Neolithic sheep individuals at 18 SNP markers linked to putatively positively selected loci in domestic sheep[3]. Argali (*Ovis ammon*) and Vignei (*Ovis vignei*) sheep were included for ancestral state inference (e.g. at SNP position OAR6_76473607.1, the ancestral state may be inferred to be C). Empty cells indicate that the allele has not been detected in an ancient individual. The modern genotype frequencies are calculated based on data from Kijas et al.[3]. Genes in putative selected regions and their functional annotations are shown within parentheses (adopted from Kijas et al.[3]. Ref/alt: reference and alternative alleles.

| **Chr** | **SNP ID** | **Ref / alt** | **Modern domestic** (*n*=749) | **Argali** (*n*=6) | **Vignei** (*n*=4) | **TEP03** | **TEP62** | **TEP83** | **ULU31** |
|---|---|---|---|---|---|---|---|---|---|
| 2 | s29378.1 | A/G | 0.96 | 1.00 | 1.00 | - | A | A | A |
| 2 | s20468.1 (*NPR2*: skeletal morph,body size) | G/A | 0.51 | 1.00 | 1.00 | A | A | - | A |
| 2 | s01865.1 | T/C | 0.96 | 1.00 | 1.00 | T | T | - | - |
| 3 | OAR3_141586525.1 | C/T | 0.67 | 0.92 | 0.50 | C | C | - | C |
| 5 | s36709.1 | A/G | 0.97 | 1.00 | 1.00 | A | A | - | - |
| 6 | s21552.1 (*FGF5*: hair variation in dogs) | G/A | 0.87 | 1.00 | 1.00 | - | G | - | G |
| 6 | OAR6_40277406.1 | G/A | 0.96 | 1.00 | 1.00 | G | G | - | G |
| 6 | OAR6_76473607.1 (*KIT*: pigmentation in cattle, pigs) | T/C | 0.59 | 0.00 | 0.00 | C | T | - | C |
| 7 | s69881.1 | A/C | 0.998 | 1.00 | 1.00 | A | A | A | A |
| 8 | OAR8_67529714.1 | G/T | 0.52 | 0.00 | 0.00 | T | T | G | - |
| 10 | OAR10_29511510.1 (*RXFP2*: horn absence) | C/T | 0.89 | 0.92 | 0.25 | C | T | - | T |
| 11 | OAR11_18701428.1 | A/G | 0.72 | 0.00 | 0.00 | G | G | - | A |
| 13 | OAR13_51852034.1 (*BMP2*: skeletal morph,body size) | G/A | 0.90 | 1.00 | 1.00 | G | A | - | - |
| 16 | OAR16_41943180.1 (*PRLR*: milk traits in cattle) | A/C | 0.63 | 1.00 | 1.00 | A | A | A | - |
| 17 | s41543.1 | C/T | 0.98 | 1.00 | 1.00 | C | C | - | - |
| 19 | s38567.1 | A/G | 0.76 | 0.00 | 0.00 | G | G | G | - |
| 25 | s03686.1 | T/C | 0.45 | 0.00 | 0.00 | T | C | - | - |
| 25 | s10489.1 | A/G | 0.87 | 0.17 | 0.62 | - | A | A | A |

**Supplementary Figures**

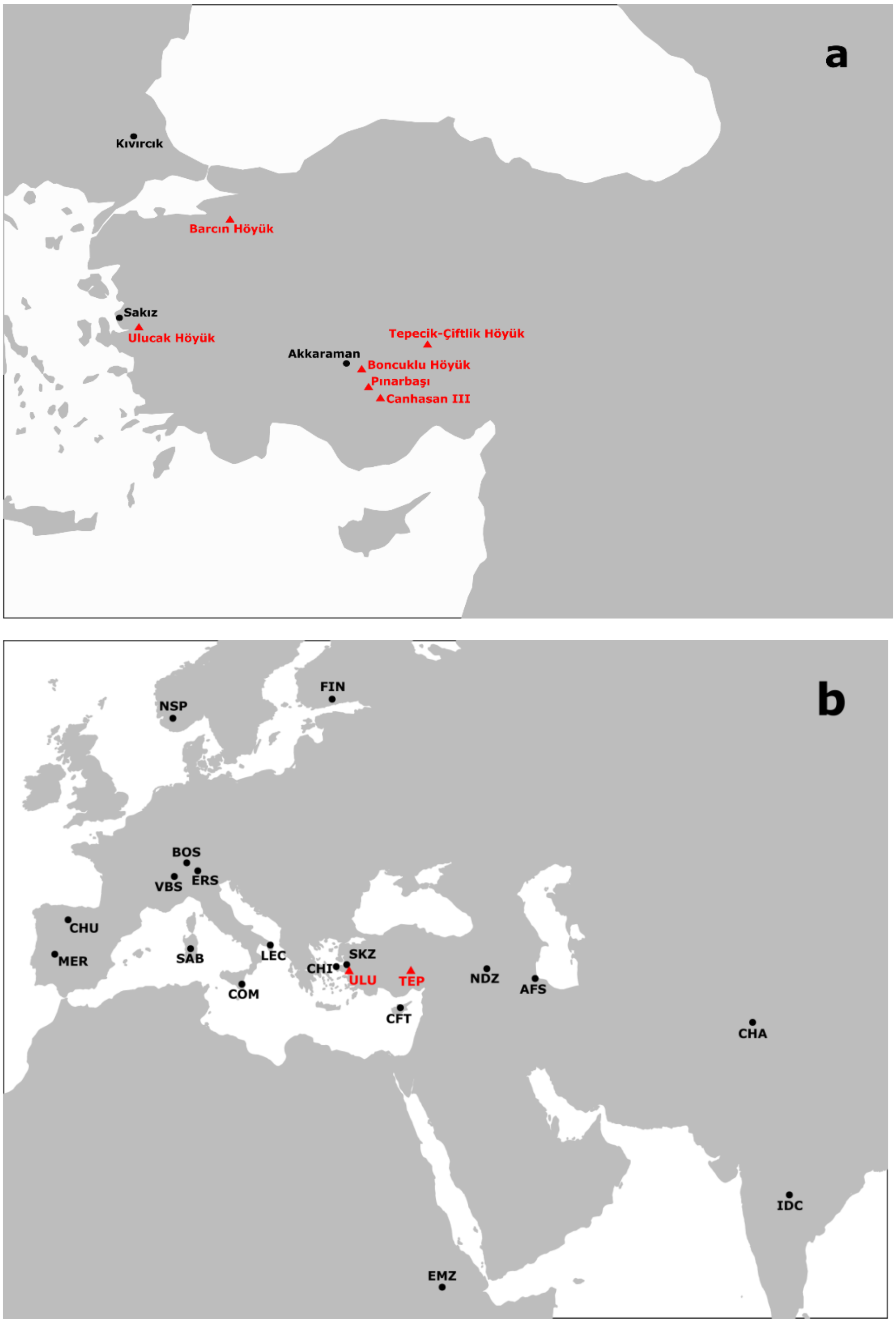


**Supplementary Figure 1.** Geographic map of modern and ancient sheep samples in this study.

(a) Geographic locations of modern breeds (in black) and ancient Anatolian sheep individuals (red) used in mitochondrial DNA analyses. (b) Geographic locations of modern domestic breeds (black) and ancient individuals (red) used in genomic analyses. For population abbreviations see Supplementary Table 5. ULU: Ulucak Höyük; TEP: Tepecik-Çiftlik Höyük.

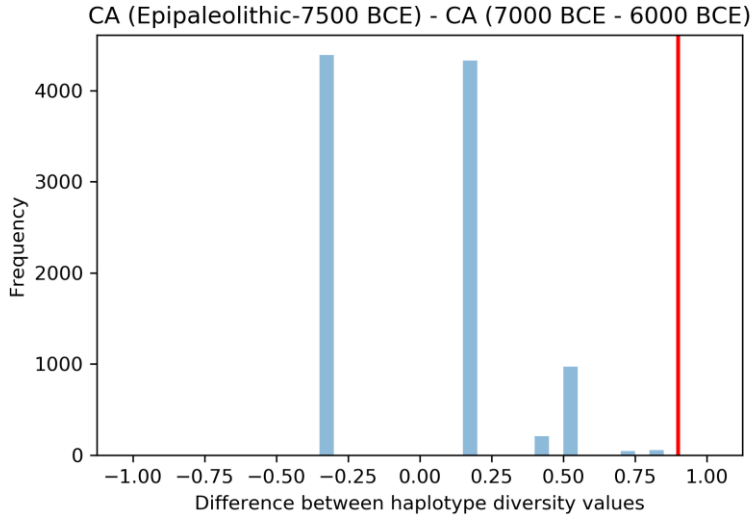


**Supplementary Figure 2.** Testing mitochondrial DNA haplotype diversity differences between different periods.

The graph shows the observed difference in central Anatolian (CA) sheep haplotype diversity (*hd*) comparing data from Epipaleolithic-7500 BCE and data from 7000-6000 BCE (red line), and the null (randomly expected) distribution of differences calculated using 10,000 permutations of group identities (light blue bars). Supplementary Table 6 shows significance values calculated using the same approach for other periods.

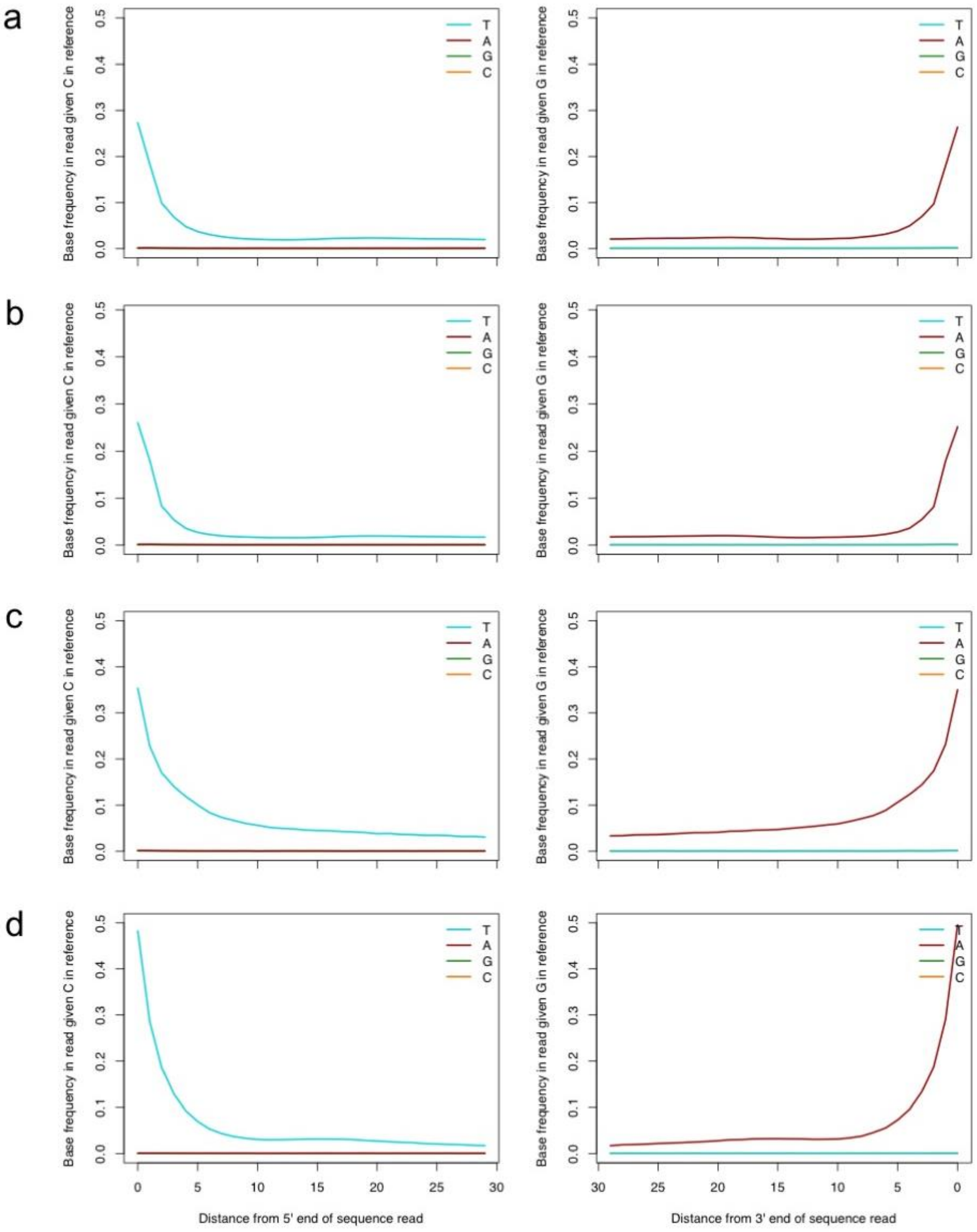


**Supplementary Figure 3.** Nucleotide mismatch patterns in Anatolian Neolithic sheep libraries used in genomic analyses.

The graphs show nucleotide mismatch patterns indicative of postmortem damage for (a) TEP03, (b) TEP62, (c) TEP83, and (d) ULU31. The y-axes indicate the frequency of mismatches between sequenced reads and the reference genome as a function of the distance (x-axis) from read 5’ ends (left panels) or 3’ ends (right panels) for each of the Anatolian Neolithic sheep used in genomic analyses.

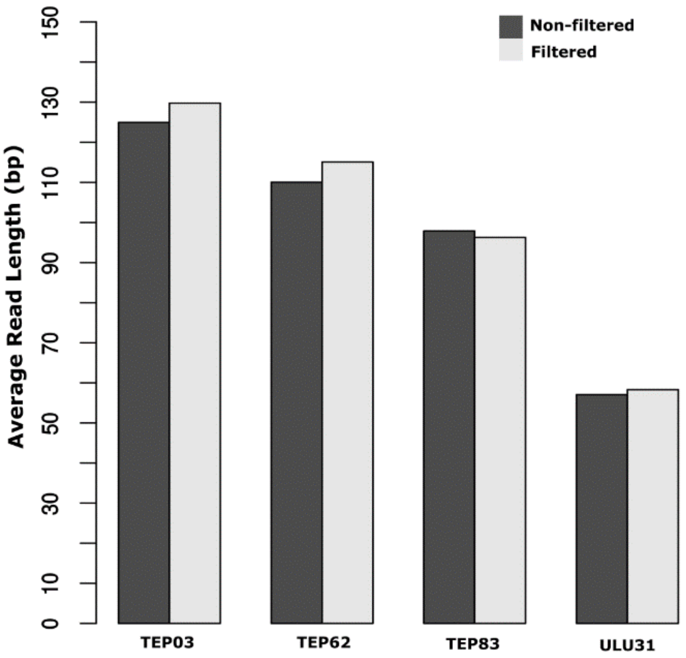


**Supplementary Figure 4.** Average read lengths of Anatolian Neolithic sheep libraries before and after filtering for postmortem damage.

Average read lengths, calculated for all reads in ANS libraries (dark grey) or only 27-53% (median 41%) of reads that pass postmortem damage (based on the C to T signature) threshold 3 set by *PMDtools*[4] (light grey). We observe no trend of shorter molecule length among filtered reads that bear postmortem-induced damage, which would be expected if long molecules represented modern DNA contamination.

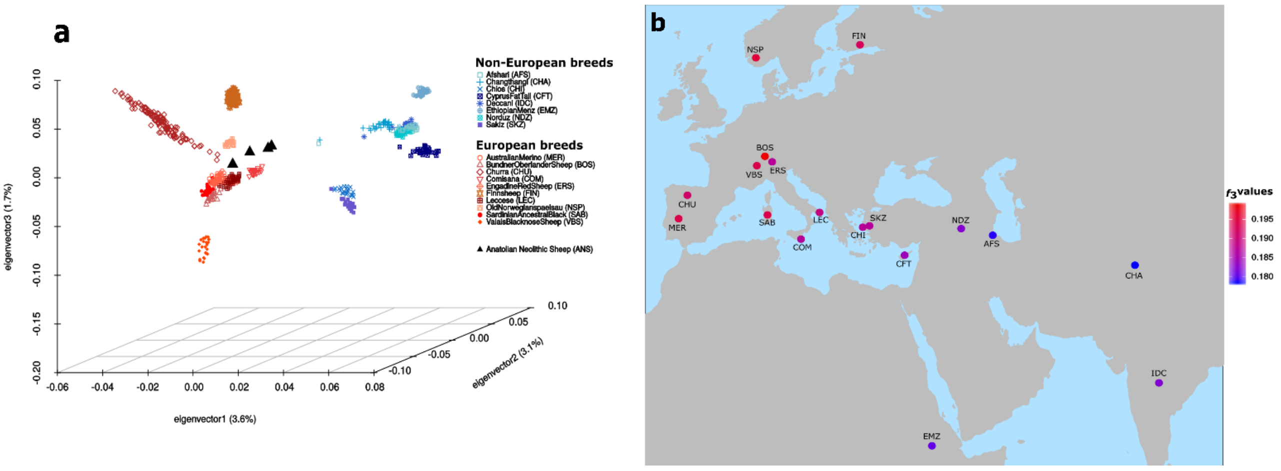


**Supplementary Figure 5.** Principal components analysis and outgroup $f_3$ plots of Anatolian Neolithic sheep and modern sheep breeds, but only using reads after filtering for postmortem damage.

The panels (a) and (b) were produced in the same way as Figures 3 and 4, respectively, but using only the 27-53% (median 41%) that pass postmortem damage threshold 3 set by *PMDtools*[4] (light grey). We observe the same patterns of affinity to European breeds in the PCA, and to north European breeds in the $f_3$ analyses, as observed using the full dataset.

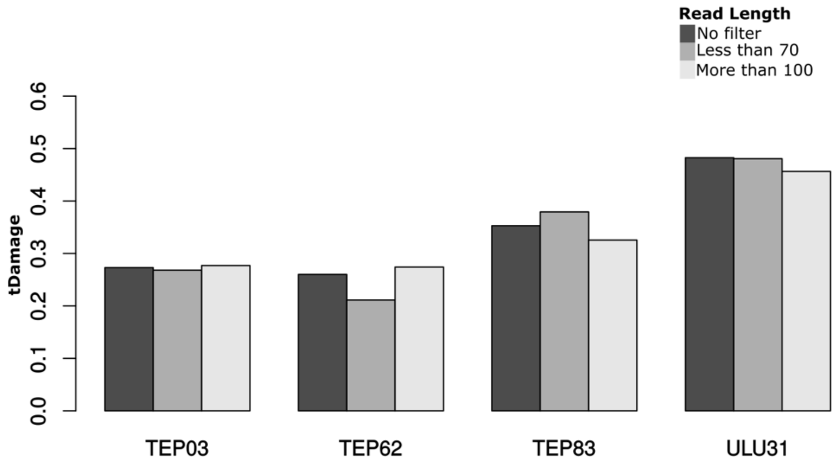


**Supplementary Figure 6.** C to T mismatch values at the 5’ ends calculated for all, short and long molecules of ANS libraries.

The barplot shows postmortem-induced C to T mismatch proportions at 5' ends across all reads ("No filter"), short reads <70 bp and long reads >100 bp, of each Anatolian Neolithic sheep (ANS) library. We observe no trend for lower C to T mismatch proportions at 5' ends for long (>100 bp) reads.

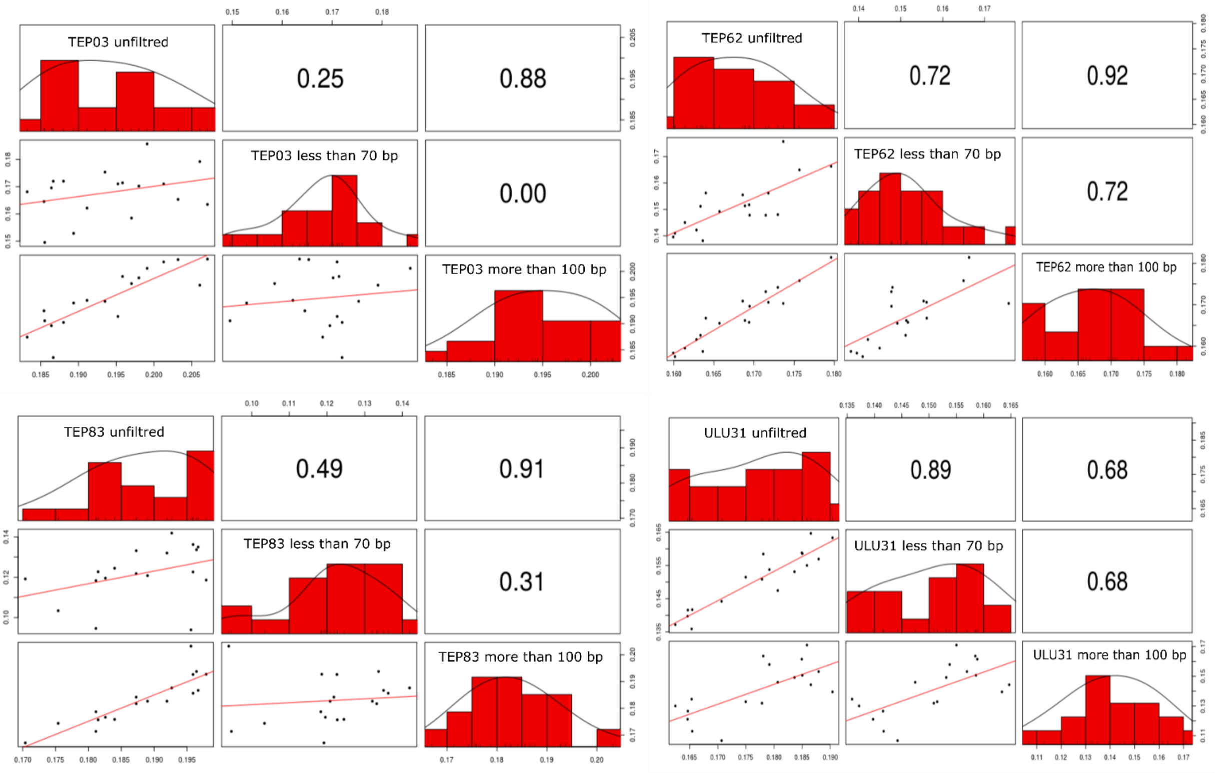


**Supplementary Figure 7.** Outgroup $f_3$-statistic distributions calculated for all, short and long molecules of ANS libraries.

The correlation plots show comparisons of outgroup $f_3$ values calculated based on different sets of molecules for each Anatolian Neolithic sheep (ANS) library. Here, $f_3$ values were calculated as $f_3$(*Argali; ANS*$_i$*, modern*), separately using genotypes called using all reads ("unfiltered"), short reads <70 bp and long reads >100 bp. In the scatter plots presented in the lower triangle, each point represents an $f_3$ value calculated between an ANS individual and a modern domestic breed, and the two axes represent $f_3$ values calculated from different molecule sets (all, short, or long). The red line represents the linear regression. The upper triangle panels show the Spearman correlation coefficients for the corresponding scatter plots. The figure was generated using the Rstudio (v. 3.5)[5] *pairs()* function with default parameters. All correlations between short and long molecules are positive, although significant (Spearman correlation test $p<0.05$) only for TEP62 and ULU31, likely due to noise in $f_3$ estimates due to low SNP numbers.

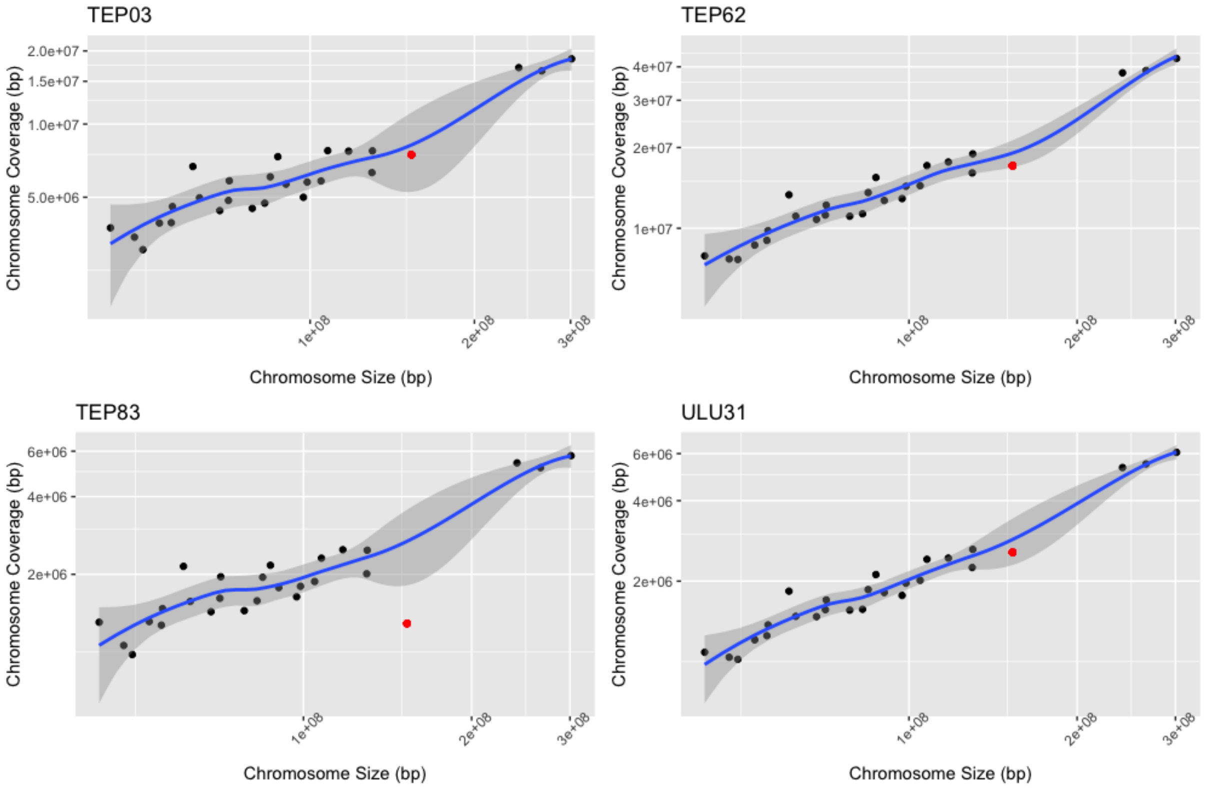


**Supplementary Figure 8.** Molecular sex estimation of the Anatolian Neolithic sheep used in genomic analyses.

Sex was estimated based on comparison of number of reads mapping to sheep chromosome X and autosomes. For each ancient individual, for all chromosomes, the number of reads (y-axis) were plotted against the size of the chromosome (x-axis). Black dots represent autosomes while chromosome X is depicted as a red dot. In each plot, 95% confidence intervals were constructed using a t-based approximation in Rstudio (v. 3.5)[5].

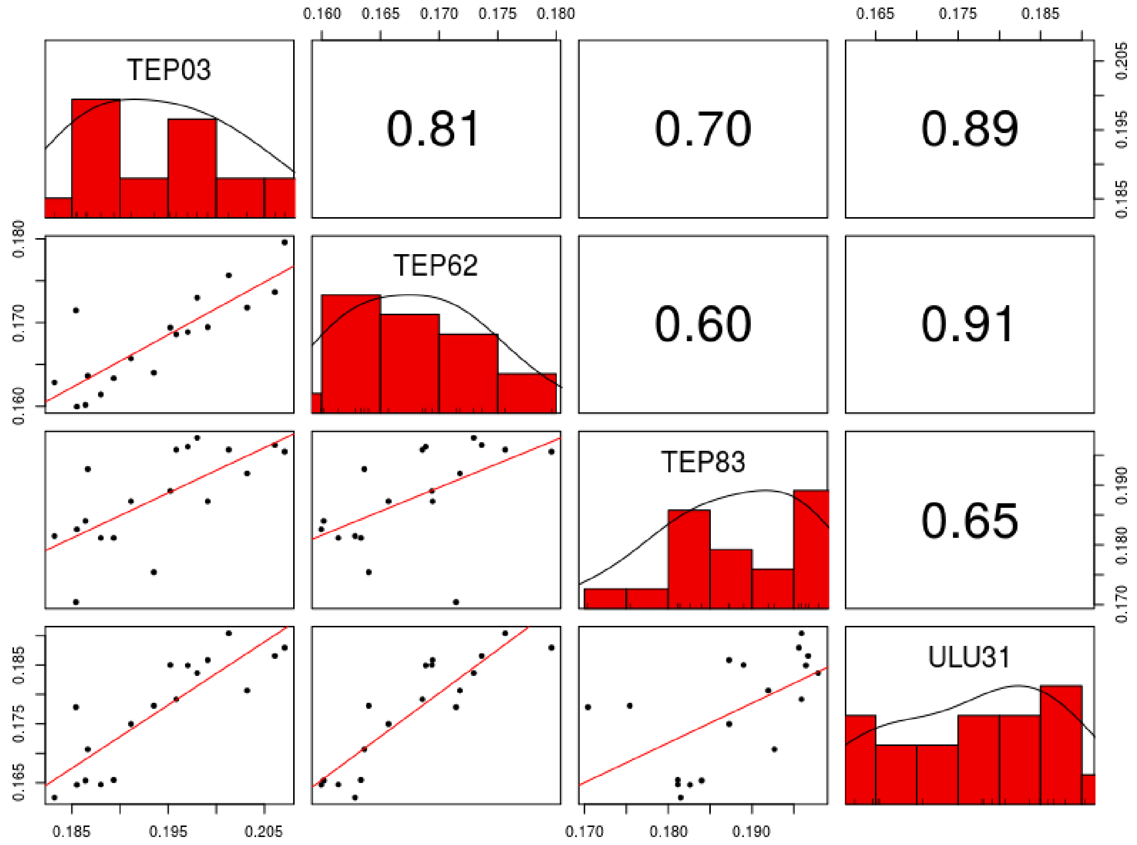


**Supplementary Figure 9.** $f_3$-statistic distributions between Anatolian Neolithic sheep (ANS) and modern sheep breeds.

In the scatter plots presented in the lower triangle, each point represents an $f_3$ value calculated between an ANS individual and a modern domestic breed, and the two axes represent $f_3$ values for two different ANS. For example, the top leftmost panel is TEP03 $f_3$ values vs TEP62 $f_3$ values, for the same modern breeds. The red line represents the linear regression. The upper triangle panels show the Spearman correlation coefficients for the corresponding scatter plots. The figure was generated using the Rstudio (v. 3.5)[5] *pairs()* function with default parameters.

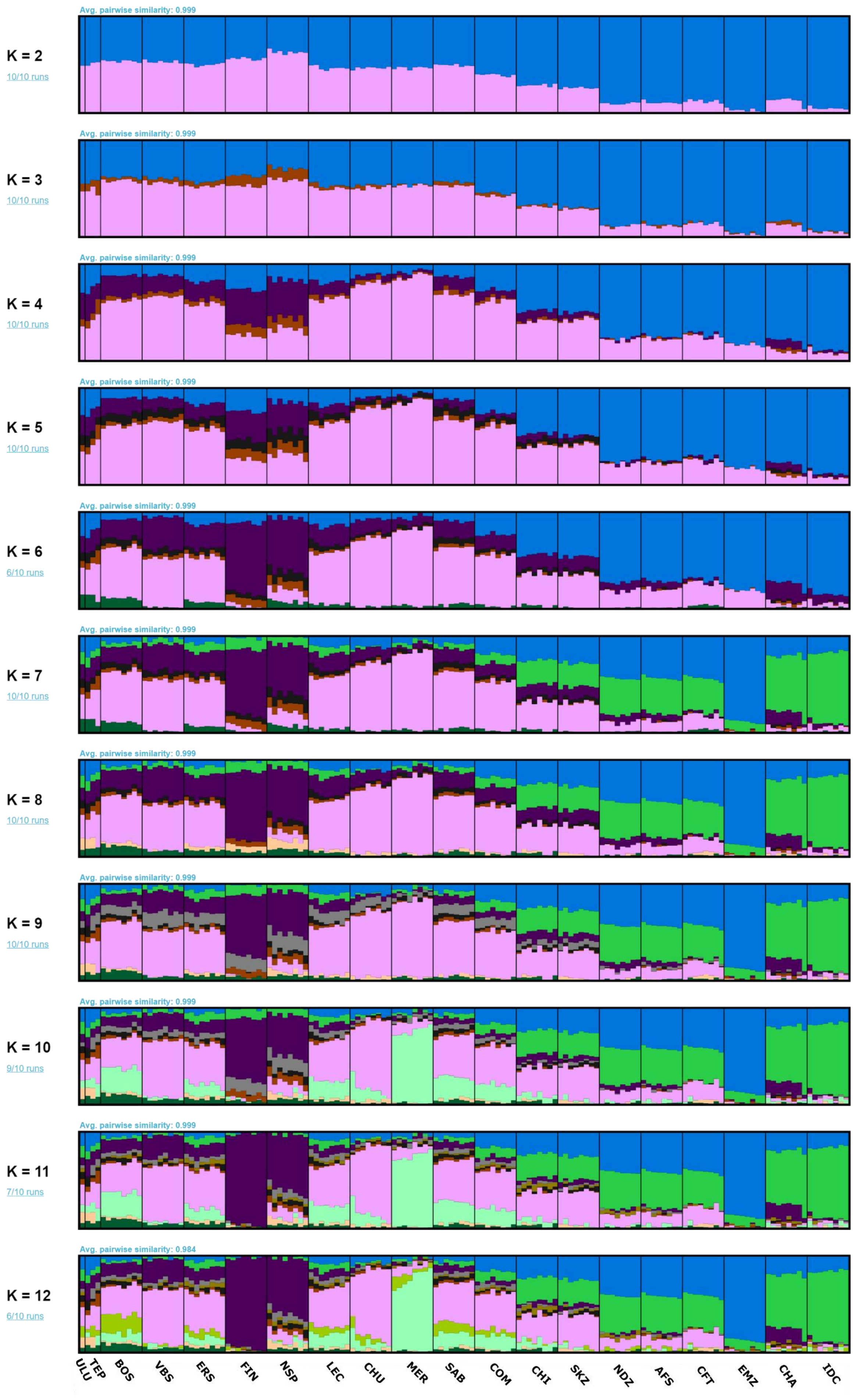

Avg. pairwise similarity: 0.999
K = 2
10/10 runs
Avg. pairwise similarity: 0.999
K = 3
10/10 runs
Avg. pairwise similarity: 0.999
K = 4
10/10 runs
Avg. pairwise similarity: 0.999
K = 5
10/10 runs
Avg. pairwise similarity: 0.999
K = 6
6/10 runs
Avg. pairwise similarity: 0.999
K = 7
10/10 runs
Avg. pairwise similarity: 0.999
K = 8
10/10 runs
Avg. pairwise similarity: 0.999
K = 9
10/10 runs
Avg. pairwise similarity: 0.999
K = 10
9/10 runs
Avg. pairwise similarity: 0.999
K = 11
7/10 runs
Avg. pairwise similarity: 0.984
K = 12
6/10 runs
ULU
TEP
BOS
VBS
ERS
FIN
NSP
LEC
CHU
MER
SAB
COM
CHI
SKZ
NDZ
AFS
CFT
EMZ
CHA
IDC

**Supplementary Figure 10.** ADMIXTURE analysis of ancient Anatolian Neolithic sheep and modern sheep breeds.

The unsupervised analysis was performed using ADMIXTURE[6]. Ten individuals were randomly chosen from each modern breed. The ANS genotypes were projected onto the calculated components. For abbreviations see Supplementary Table 7. The components were ordered and colored using the *Pong* software[7].

**Supplementary Material and Methods**

**Sample informations and archaeological contexts**

**Pınarbaşı** (by Douglas Baird)

Pınarbaşı is located on the eastern edge of the south-west Konya basin, 24.5 kms south-east of Çatalhöyük and 30 kms south-east of Boncuklu, on the tip of a limestone ridge projecting from the foot of the Karadağ volcanic massive. The site was excavated under the direction of Prof. Trevor Watkins of Edinburgh University in 1994 and 1995 and by Professor Douglas Baird of University of Liverpool in an excavation and study project running 2003-2006. The site runs from a series of cliffs on the western edge of the limestone ridge, over the slopes in front of the ridge and onto a small mound projecting into the the Hotamış lake basin. The zooarchaeological studies were carried out by Dr Denise Carruthers and Dr Caroline Middleton.

There are 4 main prehistoric periods of occupation at the site, 3 of which have provided samples for the present study. The earliest occupation so far detected at Pınarbaşı is found only in a rock-shelter next to the cliffs and is dated by C14 to the Late Glacial/Epipalaeolithic from c. 14000-11000 cal BC[8] and was excavated in Area B[8]. The second period represented at the site dates from c. 9600-7800 cal BC in the early Holocene[9] and is found only on the mound projecting into the Hotamış basin and was excavated in Areas A, C and D[9,10]. The third main period, a Late Neolithic occupation, dating c. 6500-6000 cal BC is found only in the rockshelter, was excavated in Area B, overlying the Epipalaeolithic deposits and was separated from them by a substantial amount of limestone debris. The fourth main prehistoric period, which has not contributed any samples to this study, is represented by Late Chalcolithic-Early Bronze Age occupation; covers the whole site and was excavated in areas A, C and D.

The Epipalaeolithic saw probably short episodes of seasonal occupation by small mobile foraging groups, certainly including the summer and autumn seasons, repeated over millennia, including burials as well as residential soujourns[8]. Fishing, fowling and hunting were key subsistence activities when this group camped at Pınarbaşı. Mammal hunting practices included a significant focus on caprines, the most common of the larger mammal species (c. 46% NISP) in the zooarchaeological assemblage, presumably available on the slopes of the Karadağ to the east of the site,and amongst amongst which the sheep:goat ratio is 3:2[8]. There is no evidence in their anatomical morphology or the C and N isotopes to suggest management of the caprines[11].

The 3 Epipalaeolithic samples in this study derive from several different contexts spanning most of the sequence of Epipalaeolithic occupation deposits. This sample includes 3 different haplotypes, including the haplotype that becomes predominant in the earliest probably domestic caprines after 7500 cal BC at Canhasan III. This haplotype was therefore locally present amongst sheep populations c. 6000 years before evidence of domestication. This allows the possibility of a local domestication event in central Anatolia and more specifically the Konya plain area.

The early Holocene occupation at the site dates from c. 9600-7800 cal BC[9]. This community was resident at the site on a multi-seasonal basis[9]. Sunken curvilinear plastered structures with wattle and daub superstructures were present along with significant numbers of burials. Together this evidence suggests long term investment in and commitment to the Pınarbaşı location as a residential area[9]. Baird has thus suggested this as a sedentarising behaviour. This community engaged in foraging practices with no evidence of cultivation and a focus on gathering local nuts/fruits, wild almond, terebinth and hackberry and the hunting of larger mammals, especially aurochs, accompanied by some fowling and fishing. Caprines were morphologically wild and were thus also regularly hunted, c. 27% NISP[9]. The sheep:goat ratio amongst the caprines was 8.5:1. As during the Epipalaeolithic sheep would have been quite locally available particularly on the foothills of the Karadağ to the east of the site. Study of the caprine isotopes indicate a diet very similar to that of the Epipalaeolithic caprines lacking C4 plants in the diet and eating plants low in $\delta^{15}N$ suggestng a diet not impacted by humans[11].

1 sample yielded sufficient aDNA for the purposes of this study, it derived from general midden deposits in the sequence in Area A, probably dating to the 8500-7800 cal BC phases of occupation of the site. This is HPG B, but intriguingly the haplotype of this group was not idnetified in the Epipalaeolithic sample, although this may be an issue of sample size, but it is the same haplotype as the single broadly contemporary Boncuklu sample. This haplotype is not seen in the probably earliest domestic sheep after 7500 cal BC, which indicates that the genetic bottlenecking associated with domestication may not have commenced before 7800 cal BC.

The third period of occupation at Pınarbaşı was located only in the rock shelter area and dated to the Late Neolithic c. 6500-6000 cal BC[12]. This was contemporary with the later levels at Çatalhöyük East. The occupations had a strongly seasonal natüre, with spring particularly well represented, but probably some autumn visits as well. Morphologically wild cattle and equids are attested in the assemblage, but the highest proportions of larger mammals are caprines, NISP 72.5%, morphologically domesticated and kept close to the occupation area judging by large quantities of perinatal remains and herbivore dung in deposits on the site[12,13]. Amongst the caprines the sheep:goat ratio is 168:1[12], showing a massive preponderance of sheep in the flocks and a majör shift compared to ratios in the earlier assemblages. The C and N isotopes also strongly indicate these caprines are herded, given their generally elevated $\delta^{15}N$ and significant proportion of individuals consuming C4 plants, with isotope signatures close to contemp Çatalhöyük[11,12].

Four samples in this study derived from these Late Neolithic occupation phases at the site. These show the dominance of HPG B haplotype that seems to predominate amongst the domesticated sheep of Neolithic central Anatolia and the addition of haplogroup D, not seen in the wild population in the area, so presumably representative of sheep introduced to the area by c. 6500 cal BC, interesting evidence of movement of sheep between communities during the Neolithic, through exchange, pastoralism and/or accompanying the movement of people into the Konya plain.

**Boncuklu Höyük** (by Douglas Baird**)**

Boncuklu is an archaeological mound in the middle of the south-west Konya basin in south central Anatolia. It is located 9.5 kms north of the site of Çatalhöyük. Excavations have been directed by Professor Douglas Baird from the University of Liverpool since 2006 and co-directed with Baird by Professor Andrew Fairbairn, University of Queensland and Dr Gokhan Mustafaoğlu, Ankara Haci Bayram Veli University, since 2011. The zooarchaeological studies were carried out for most of this period by Dr Louise Martin and Dr Caroline Middleton. Evidence for occupation spans 8300-7600 cal BC. The site contains a series of sub-oval mudbrick structures some with sub-floor burials, as well as burials in open areas. Buildings have standard domestic features and notably the space is divided into ‘clean’ and ‘dirty’ areas. This evidence comnbined with evidence for occupation of the site in all seasons over a prolonged period strongly suggests an early sedentary community[9]. Adoption of cultivation by indigenous foragers[14] is suggested at Boncuklu and likely by 8300 cal BC[10]. Cultivated wheat and legumes probably made a limited contribution to the diet. Cultivated plants may have been introduced through interactions with areas to the south and east in the 9th millennium cal BC. Morphologically wild cattle - aurochs, and probably wild, boar represent the main hunted species. In contrast to Pınarbaşı caprines represent a very small element of the faunal NISP, c. 4 %, with a sheep:goat ratio of c 1:1.33[9].

This is not surprising as the closest wild caprine natural habitats would be the Bozdağ c. min 15 kms to the north. However, C and N isotopes, with some individuals with elevated $\delta^{15}N$ suggest some human effect on caprine diet[11] and along with herbivore dung on the site[13] hint at early management. In that context it is notable that the single aDNA sample from the site is the same haplogroup/haplotype as broadly contemporary Pınarbaşı, unlike the somewhat later probably early domestic caprines, with little evidence of the commencement of genetic bottle necking indicated by the predominance of the haplotype seen at Canhasan III and Pınarbaşı Late Neolithic. This may well point to management definitively preceding the development of the genetic domestication process. In addition it is interesting that a haplotype that was apparently not uncommon and widespread in the 9th millennium cal BC Konya plain was not caught up in the bottle neck of domestication.

**Canhasan III** (by Douglas Baird)

Canhasan III is located on the Karaman alluvial fan south-east of the volcanic massif of Karadağ in the Konya Plain and c. 90 kms South-east of Çatalhöyük and was excavated in 1969-70 by David French[15], and is dated to c. 7400–7100 cal BC[16]. At the request of Dr French, Prof. Douglas Baird is overseeing study and analysis of material and information from the site to more substantive publication. French excavated a 20x30 m area by surface scraping in which he exposed an extensive dense set of abutting rectanglar structures similar to sites like Aşıklı and Çatalhöyük. He also excavated in a much smaller area, in square 49L, a deep sounding to 6.75 m which stopped at the water table. The plant remains indicate the exploitation of wild species as well as cultivated cereals. The animal remains, which are subject to ongoing study, indicate an important presence of cattle and caprines. The domestication status of these species is unclear[17]. A recent study of caprine diets through C and N isotopes by Middleton[11] strongly indicated that a high proportion of caprines consumed C4 plants and most individuals showied high $\delta^{15}N$, like the later Çatalhöyük East morphologically domestic caprines, were thus probably herded.

The 3 samples wihtin this aDNA study were from both the wider scraped area and the deep sounding. They are all of the HPG B and the haplotype that predominates in slightly later domestic sheep flocks and this supports the evidence of C and N istopes that the Canhasan III sheep (and probably therefore goats) were herded, with high degrees of management and, in addition, that this had resulted in genetic domestication. This evidence also ties down the appearance of genetically domestic animals on the Konya Plain to the period between c. 7800 and 7300 cal BC.

**Tepecik-Çiftlik** (by Erhan Bıçakçı and Yasin Gökhan Çakan)

Tepecik-Çiftlik mound is located in the Çiftlik district of Niğde province which resides in the Volcanic Cappadocia region of the Central Anatolian Plateau. The 6-hectare mound has a height of 9,60 meters above the plains it is located. Although the mound was excavated 7,30 meters below the ground, the rock soil has not been reached yet[18]. Excavations carried out in the mound since the year 2000 pointed out that from the end of the 8th millennium BCE to the beginning of the 6th millennium BCE the settlement was occupied by an agriculturalist community. Alongside the farming, the community continued hunting and gathering the rich resources of the region. Significant changes in the architecture and settlement layout could be observed during the 1200-year period that straddled from the early Pottery Neolithic towards the end of early Chalcolithic[19]. Almost every level has a different layout. Although the buildings in different levels were built with similar techniques, they differ from each other in terms of space perception and settlement pattern. Sheep and goat were the highly dominant animal species in the settlement. They were followed by cattle, equids and red deer, respectively. Additionally, the other species detected among the animal remains were roe deer, fallow deer, canids, rabbit, bear, rodents, pig and some bird species. Domesticated sheep, goat and cattle were observed after the first half of the 7th millennium BCE. However, hunting wild animals was also part of the subsistence[20,21]. The oldest known pottery in the Volcanic Cappadocia region is found in Tepecik-Çiftlik[22]. The continous stratification in the mound permits to follow both technological and typological development from the first time the pottery appeared  to the end of the early Chalcolithic period.  The obsidian production areas within the settlement emphasizes the significance of processing and the use of this raw material for the community[23]. Almost all types of the arrow and spearheads known from the contemporaneous Central Anatolian settlements are also found in Tepecik-Çiftlik. It is known that the obsidian blades and cores produced in Göllüdağ workshops made their way to the settlements in far away regions such as the Levant, Northern Syria and Cyprus since the earliest phases of the Neolithic period[24]. Networks established between the obsidian workshops and settlements could be regarded as the most concrete indication of the mutual transfer of knowledge and technology between these regions. It should be noted that the Tepecik-Çiftlik people were not ignorant of the developments taking place in the workshops immediately next to the settlement[25]. Archaeological discoveries in Tepecik-Çiftlik indicate a community that could make good use of the natural resources around it and that retains close relationships with the nearby as well as the distant regions.

**Ulucak Höyük** (by Özlem Çevik)

Ulucak Höyük, lies 25 km east of İzmir in West Central Turkey, is yet to be a key site in the region. The Neolithic occupation at the site is designated by Levels VI through IV, and is dated between 6850 calBC and 5670 calBC[26]. A full-fledged agricultural economy with cereals and pulses and four-tiered herding system including sheep, goat, cattle and pig has been attested starting from the basal levels onwards[27]. The earliest occupational level is devoid of pottery and any other clay objects while sporadic occurrence of Melian obsidian in this level suggests contacts with the Aegean world was already established[28,29]. Pottery was adopted after 6600/6500BC although clay images including figurines and seals became an integral part of the material culture towards the end of the 7$^{th}$ millennium BC[30,31]. The initial settlement (Level VI: 6850-6500 calBC) has so far been represented by two rectangular buildings with mud slab walls (Building 42 and 43) flanked by open spaces with fire installations.  The floors and possibly the walls of these buildings were lime plastered and red painted. As the buildings appear to have been deliberately left clean, sheep bone samples were taken around the fire installations. Level V (6500-6000 calBC) with its five sub phases (a-e) is represented by post-framed and mud-slab buildings while both substantial mud-brick buildings on stone foundations and post-framed buildings are found in Level IV (6000-5670 calBC).

**Barcın Höyük** (by Rana Özbal and Fokke Gerritsen)

Barcın Höyük, a seventh millennium Neolithic site, is located in the Yenişehir Plain in the province of Bursa in northwestern Turkey[32]. Excavations here, directed by the Netherlands institute in Turkey, yielded an uninterrupted Neolithic sequence from 6600 cal BC through 6000 cal BC. The lowest levels of the settlement represent the initial occupation by pioneer farmers who arrived in this region presumably from Central Anatolia[33]. These people brought with them domesticated stocks of plants and animals in a conscious effort to settle and colonize these regions[34,35]. It is possible that small numbers of foragers inhabited the region but the pioneers at Barcın Höyük maintained a fully Neolithic lifestyle[36]. They lived in rectangular timber structures organized in rows and surrounded by courtyard spaces[37]. The courtyards were used for domestic activities and functioned as cemeteries especially for the adult population. Infants were buried beneath the floors of the houses while children were placed just outside the houses in the annexes or veranda spaces. The earliest inhabitants were accomplished potters though they did not initially rely on this technology for cooking or for storage and instead make use of indirect cooking techniques[38]. Pottery recipes were perfected through time; by 6500 BC the pots show expert burnishing and new tempers types were added to strengthen the vessels[37]. Domesticated cereals and pulses comprised the majority of the plant foods consumed although hazelnuts were also exploited[39]. With regards to domesticated animals, cattle and sheep dominate the assemblage[40]. Domestic goats too are found though in small numbers. Evidence for hunting and hunted animals is limited at Barcın Höyük although the faunal record included small numbers of wild boars, deer, hare. Dairying presumably from cattle but potentially also sheep was part of the diet from the earliest levels onwards[35].

### Laboratory analyses

#### Sample preparation and DNA extraction

DNA extractions and library preparations were conducted in a dedicated ancient DNA laboratory at the Middle East Technical University. Prior to DNA extraction, outer surfaces of bone samples were cleaned by sandblasting with a cutting disk attached to a Dremel tool. A new cutting disk was used for each sample. DNA extractions were performed following Dabney et al.[41]. A small piece of bone was cut out of cleaned bone and then ground to fine powder with a mortar and pestle. 120-200 mg of bone powder was weighed and placed in a 2 ml screw-top tube. Briefly, 1 ml extraction buffer (0.45 M EDTA and 0.25 mg/ml ProteinaseK) was added onto bone powder and incubated at 37 ºC for 18 hrs in a rotator incubator After incubation, tubes were centrifuged and supernatants were transferred into tubes containing 13 ml binding buffer (M Guanidine Hydrochloride, 40% (vol/vol) Isopropanol, 0.05% Tween-20, 90mM Sodium Acetate). This mixture was filtered through Qiagen Minelute spin columns to bind and wash DNA which was followed by two consecutive elutions with 50 µl Qiagen EB buffer.

#### PCR amplification and sequencing of mtDNA

Amplification of aDNA samples for 144 bp mtDNA control region (CR) was performed in a volume of 20 µL containing, 1X Taq buffer, 2 mM $MgCl_2$, 0.25mM dNTPs, 15 pmol of each primer, 2 ng BSA and 3 unit of Taq DNA polymerase. Thermocycling conditions were as follows: initial denaturation at 94°C for 10 minutes, followed by 60 cycles of denaturation at 94°C for 30 seconds, annealing at 53°C for 45 seconds, extension at 72°C for 45 seconds and a final extension at 72°C for 5 minutes. Sanger sequencing of the PCR products were carried out by Refgen Inc. The 144 bp long fragment of mtDNA corresponding to the positions 15391–15534 on the reference mitogenome (AF010406) was sequenced from 74 ancient samples by using primer pairs from Cai et al.[42]. mtDNA haplogroups (from A to E) of samples were assigned according to the identity of nucleotides on HPG determining positions with respect to the reference AF010406 sequence (TableS3).

### Population genetic analyses of mtDNA data

#### Haplogroup assignment and frequency comparisons

The 144 bp sequences were assigned to haplogroups as described in the main text. For testing the hypothesis that haplogroup B haplotype frequencies changed between periods, we used the Fisher's exact test (two-sided) using the *R* (v. 3.5)[5] *fisher.test* function.

#### Haplogroup and haplotype diversity

Haplogroup diversity was estimated using the Shannon Diversity Index (*H*). In addition, nucleotide and haplotype diversity values were calculated using *DnaSP* v6[2]. The statistical significance of the difference between the diversity values of groups was determined by a two-sided permutation test, performed using custom *Python* code. Specifically, for each pairwise comparison, pooled samples were randomly allocated into two groups in accordance with the original sample sizes and diversity values were calculated. This procedure was repeated 10,000 times and a null distribution of possible values were obtained. The difference between groups was assumed significant if the frequency of the observed value was less than 5% within this null distribution.

#### Whole genome shotgun sequencing, in-solution SNP capture and data preprocessing

Double-stranded Illumina libraries were built for 36 sheep samples (Table S8) following Meyer and Kircher[43] and sequenced on the HiSeq 4000 platform. In order to increase coverage, libraries with >1% endogenous sheep DNA (TP001, TP062, TP83, UH026, UH031) were enriched for 20,000 SNPs via in-solution capture hybridization using custom designed MyBaits probes (Arbor Biosciences Inc.) following the manufacturer's instructions (see below for SNP selection and probe design). In-solution hybridization capture was carried out following manufacturers instructions. Hybridizations reactions were incubated at 55ºC for 24 hours and the captured libraries were amplified using Herculase II Fusion Polymerase (Agilent Technologies) for 15-19 cycles. Enriched libraries were purified using AMPure XP beads and then quantified by Agilent 2100 Bioanalyzer. Purified libraries were pooled in equimolar concentrations to reach a final concentration of 10nM and then sequenced on the HiSeq X platform.

#### SNP selection and capture probe design

For in-solution-hybridization capture, 20,000 SNPs were selected from Illumina's OvineSNP50 Genotyping Beadchip SNPs[3]. Four 60 nucleotide- long probes were designed per SNP, summing up to a total of 80K probes for 20K SNPs. Two of the probes, carrying either the reference or the alternative allele, were centered on the SNP, while the other two (carrying the reference sequence) were located at each side of the targeted SNP, following the design by Haak et al.[44]. The 20,000 SNPs included all 8850 transversions present in the Illumina OvineSNP50 Beadchip variant set, 1237 chromosome X transitions, 3 mtDNA transitions, and 521 putatively functional SNPs [based on Kijas et al.[3], Moradi et al.[45], Heaton et al.[46], Noce et al.[47]]. The remaining 9389 SNPs were randomly selected among OvineSNP50 transitions. The designed probes were produced by Arbor Biosciences Inc.

#### Data preprocessing

We processed the data following Kılınç et al.[48] and Günther et al.[49] . We removed the residual adapter sequences in FASTQ files and merged the paired-end sequencing reads using MergeReadsFastQ_cc.py[50]. We mapped the merged reads to the sheep reference genome (Oar_v3.1) using *BWA* (v. 0.7.12)[51] *aln* algorithm, using the parameters: '*-n 0.01 -o 2 –l 16500*'. After merging all libraries from the same individual using *samtools*[52] merge function, we removed the PCR duplicates by using FilterUniqueSAMCons.py[50]. We filtered out reads shorter than 35 base pairs. Additionally, reads with greater than 10% mismatches to the sheep reference genome were removed.

#### Authentication of ancient sequences

In order to verify the authenticity of the genomic data we assessed post-mortem damage patterns specific to ancient DNA and not expected in modern DNA data. Due to deamination of cytosines at the 5' ends of the broken DNA fragments creating uracils or thymines, we expect to observe frequent cytosine to thymine transitions when comparing ancient reads with the reference genome sequence[53]. The software *PMDtools* was used to evaluate nucleotide misincorporation patterns at the first 60 positions at 5' and 3' ends of the reads[4]. The frequencies of C to T transitions at the 5' ends ranged between 42% to 20% (Supplementary Figure 3), supporting the authenticity of the bulk of the sequences. We filtered reads for postmortem damage using *PMDtools*[4] with the *'--threshold 3'* parameter. We then compared the PMD-bearing reads with the unfiltered read set, with respect to read lengths, and population genetic characteristics, namely PCA and outgroup $f_3$-value distributions. Additionally, we prepared two different read sets for each Anatolian Neolithic sheep individuals. We generated two different BAM files for each ANS individual by choosing read lengths as less than 70 bp and more than 100 bp using "awk script" commands on Linux terminal. We conducted f3 analysis by following same procedure with supplementary methods of f3-statistics. Finally, f3 correlations of each individual were compared (unfiltred, less than 70 bp and more than 100 bp of one individual).

**Read trimming and SNP calling**

BAM files were trimmed for 10 bp from both ends to avoid confounding of post-mortem damage during genotype calling process using trimBam[54] We used the Illumina OvineSNP50 Beadchip variant set[3] for genotype calling using *SAMtools* (v. 1.3)[52] *mpileup* program with the parameters: '*-q 30 -Q 30*'. We randomly pseudohaploidised the genotypes using custom *python* code following Kılınç et al.[48].

**Molecular sex determination**

The molecular sex of the four individuals was estimated by comparing the number of reads mapping to autosomes and chromosome X. The number of mapped reads were normalized by the length of chromosome. Considering males have a single copy of chromosome X, they are expected to have 50% of reads mapping to chromosome X compared to autosomes, unlike females, which were expected to have comparable number of reads mapping to chromosome X and autosomes. Graphs were constructed using *ggplot2*[55] in *R* (v. 3.5)[5], with the *stat_smooth()* function to construct a regression curve with method *loess* (due to less than 1000 observations) and a span value of 0.80 (fraction of points used to fit the curve) to control the amount of smoothing. Confidence intervals (95%) were constructed using a *t*-based approximation.

**<u>Population genetic analyses</u>**

**Merging ancient and modern sheep genotypes**

As described in the main text, we chose 18 modern sheep breeds from the 74 worldwide breeds included the Kijas et al.[3] SNP chip dataset (Supplementary Table 2). As the modern data set had been generated using oviAri1.v1 reference genome coordinates, these were mapped to oviAri3 coordinates using the UCSC Genome Browser *liftover* tool[56]. Out of 49,034 SNPs in the SNP chip dataset, 40,225 could be mapped to the oviAri3 genome version and were used in subsequent analyses. We merged modern and ancient genotype dataset using the *PLINK* (v. 1.9) software[57] '*-mergelist*' command, to produce a dataset with eighteen modern populations and four ANS samples.

**Principal component analysis (PCA)**

Eigenvalues of each PCA were calculated using the "*smartpca*" program of *EIGENSOFT* (v. 7.2) software[58], using all eighteen modern domesticated breeds' genotypes. The four ancient samples' genotypes were then projected onto the space described by these eigenvalues using *EIGENSOFT*'s "*lsqproject:YES*" parameter. All graphs were drawn using the "*scatterplot3d*" package[59] and *plot*() function of Rstudio (v. 3.5) software[5].

***D*-statistics**

We compared multiple combinations of modern and ancient populations to address questions about relationships among populations and thus infer demographic history. *D*-statistics were conducted using the *qpDstat* program of *AdmixTools* (v. 5.1) software[60]. Modern, ancient and outgroup datasets were merged using the "mergelist" command; extra chromosomes of sheep that were not recognized by *PLINK*[57] were excluded by "*exclude*" command. PED, PEDIND and MAP file formats were converted into *EIGENSOFT* file formats[60], namely GENO, IND and SNP file formats, before *D*-statistics tests were performed. We chose the Argali sheep (*Ovis ammon*) as outgroup in this study, after confirming that Argali sheep was equally distant to all modern sheep breeds in *D*-tests using the goat as outgroup: *D(goat, Argali; modern1, modern2)*. The *p*-values were calculated from *Z* scores based on the block jackknife procedure[60]. In order to control for the false discovery rate, we performed multiple testing correction using the "*p.adjust*" function in *R* (v.3.5)[5], using the Benjamini-Hochberg method[61] , for each batch of tests performed.

#### $f_3$-statistics

For calculating outgroup $f_3$-statistics, modern and ancient dataset were merged with and converted into *EIGENSTRAT* file format using the same procedure applied in *D*-statistics. First, the test population was Anatolian Neolithic sheep individuals separately, and they were compared with modern sheep breeds populations one by one, calculating $f_3$(*Argali_individual; ANS_individual, modern*). The significance of the positive correlation between the resulting $f_3$ values for each pair of ancient individuals was calculated using the Spearman correlation test in *Rstudio* (v.3.3)[5]. After confirming high correlations, all four ancient individuals were treated as a single ANS population, and we performed tests as $f_3$(*Argali; ANS, modern*). The significance of higher $f_3$ values in European-ANS comparisons vs. non-European-ANS comparisons was calculated using the Mann-Whitney U-test in *Rstudio* (v.3.5)[5].

In order to prepare ancient and modern data for admixture $f_3$-statistics, the same procedure that was applied in outgroup $f_3$-statistics was followed, except that instead of an outgroup, we calculated $f_3$-statistics as $f_3$(*modern1; ANS, modern2*), where *ANS* represents the genotype of all four Anatolian Neolithic sheep combined, and *modern1* and *modern2* represent the genotypes of modern south European and modern Asian breeds, respectively. We conducted admixture $f_3$-statistics using *Admixtools* (v. 5.1) software using the *'inbreed: YES'* parameter[60].

#### ADMIXTURE analysis

To estimate ancestral components in Anatolian Neolithic sheep and in the 18 modern sheep breeds, we performed unsupervised clustering analysis using the *ADMIXTURE* software[6]. We first filtered modern sheep breed dataset for linkage disequilibrium using *PLINK* [57], with parameters '*--indep-pairwise 50 5 0.05*'. All SNPs filtered (*n*=2265, 19.5%)  from the modern dataset were also excluded from ancient individuals using *PLINK* [57]. We then carried out cluster analysis for modern sheep breeds using between *K*=2 and *K*=12 components. For every *K* value, we generated 10 replicate runs, repeating the analysis using different random seed numbers. Finally, we used the "projection" function of *ADMIXTURE* with random different seed and again performed 10 runs following, to calculate Anatolian Neolithic sheep individuals' ancestral components based on those calculated from modern breeds. We used the *Pong* [7] program for visualization. We randomly chose 10 individuals for each modern sheep breed to improve visualization.

#### SNPs at putatively selected loci

Kijas et al.[3] had identified 31 putatively selected loci using an *Fst* scan across global sheep breeds, comprising the most extremely differentiated 0.1% of genome-wide markers. The highest ranking 31 SNPs for each locus was represented on our 20K capture probe set, and eighteen of these SNPs were present in at least one of the four ancient sheep samples' libraries. We compared these ancient sheep genotypes with those of 18 populations of modern domestic sheep breeds (used in the main analyses), as well as with two outgroups, Argali and Vignei, to identify the ancestral state. Data for both the outgroups and the modern sheep were retrieved from the Kijas et al.[3] dataset.
